# *Lactiplantibacillus plantarum* uses ecologically relevant, exogenous quinones for extracellular electron transfer

**DOI:** 10.1101/2023.03.13.532228

**Authors:** Eric T. Stevens, Wannes Van Beeck, Benjamin Blackburn, Sara Tejedor-Sanz, Alycia R. M. Rasmussen, Emily Mevers, Caroline M. Ajo-Franklin, Maria L. Marco

## Abstract

Extracellular electron transfer (EET) is a metabolic process that frequently uses quinones to couple intracellular redox reactions with extracellular electron acceptors. The physiological relevance of this metabolism for microorganisms that are capable of EET, but unable to synthesize their own quinones, remains to be determined. To address this question, we investigated quinone utilization by *Lactiplantibacillus plantarum,* a microorganism required for food fermentations, performs EET, and is also a quinone auxotroph*. L. plantarum* selectively used 1,4-dihydroxy-2-naphthoic acid (DHNA), 2-amino-3-carboxy-1,4-naphthoquinone (ACNQ), 1,4-naphthoquinone, and menadione for EET reduction of insoluble iron (ferrihydrite). However, those quinones used for EET also inhibited *L. plantarum* growth in non-aerated conditions. Transcriptomic analysis showed that DHNA induced oxidative stress in *L. plantarum* and this was alleviated by the inclusion of an electron acceptor, soluble ferric ammonium citrate (FeAC), in the laboratory culture medium. The presence of DHNA and FeAC during growth also induced *L. plantarum* EET metabolism, although activity was still dependent on the presence of exogenous electron shuttles. To determine whether quinone-producing bacteria frequently found together with *L. plantarum* in food fermentations could be a source of those electron shuttles, *L. plantarum* EET was measured after incubation with *Lactococcus lactis* and *Leuconostoc mesenteroides.* Quinone-producing *L. lactis,* but not a quinone-deficient *L. lactis* Δ*menC* mutant, increased *L. plantarum* ferrihydrite reduction and medium acidification through an EET-dependent mechanism. *L. plantarum* EET was also stimulated by *L. mesenteroides*, and this resulted in greater environmental acidification and transient increases in *L. plantarum* growth. Overall, our findings revealed that *L. plantarum* overcomes the toxic effects of exogenous quinones to use those compounds, including those made by related bacteria, for EET-conferred, ecological advantages during the early stages of food fermentations.

## Introduction

Oxidation-reduction (redox) reactions dictate the flow of electrons in many enzymatic reactions. Accordingly, these reactions are facilitated by an abundance of shuttling compounds used by bacteria to transport electrons within and outside of the cell (Zerfaß et al., 2019). Quinones are one such class of diverse, redox-active molecules with over 1,000 structural forms (Taylor and Francis, 2022). Quinones serve numerous functions in cells in all domains of life including in electron transport chains used for respiration energy conservation metabolism, extracellular protein folding, and pyrimidine synthesis (Franza and Gaudu, 2022). Conversely, these compounds can also cause cellular damage. Quinones reduce oxygen to reactive oxygen species (ROS) that result in DNA damage and membrane lipid peroxidation (Goel et al., 2020). Exogenous quinones can also inhibit respiration by competing with native quinones produced by the cell (Nagata et al., 2010). The conflicting impacts of quinones on energy conservation pathways illustrate the complexity of their effects. Furthermore, their importance for microorganisms that do not rely on respiratory metabolism remains unclear.

Quinones are frequently utilized for extracellular electron transfer (EET), a metabolic process used by certain microbes that couples intracellular redox reactions to extracellular electron acceptors or donors like redox-active metals or electrodes (Doyle and Marsili, 2018; Lovley and Holmes, 2022). EET metabolism can be categorized based on the direction of electron flow to and from the electro-active microorganism and how electron transfer is completed (Lovley and Holmes, 2022; Zhao et al., 2020). The reduction of long-range electron acceptors through mediated electron transfer (MET) occurs with endogenous, membrane-associated quinones used as intracellular electron shuttles and with environmental or secreted quinones functioning as extracellular electron shuttles (Light et al., 2018; Mevers et al., 2019). These EET pathways have thus far been best characterized for the Gram-negative, mineral-respiring bacteria species of *Shewanella* and *Geobacter* (Zou et al., 2021), although the number of EET active microorganisms is now understood to extend to all three domains of life (Logan et al., 2019).

We recently showed that *Lactiplantibacillus plantarum* performs EET by a blended metabolism combining features of respiration and fermentation (Tejedor-Sanz et al., 2022b). *L. plantarum* is member of the lactic acid bacteria (LAB), a group of Gram-positive bacteria in the Bacillota (formerly Firmicutes) phylum that share metabolic and physiologic characteristics and named for their production of lactic acid from fermentative energy conservation metabolism (Gänzle, 2015). *L. plantarum* EET partially resembles respiration because EET increases intracellular NAD^+^/NADH ratios, but fermentation energy conversation metabolism with substrate-level phosphorylation is still used for ATP generation. *L. plantarum* EET results in a shortened lag phase, increased fermentation flux through organic acid production, and greater environmental acidification (Tejedor-Sanz et al., 2022b). The *L. plantarum* EET pathway is present in other LAB and similar to the flavin-mediated FLEET system encoded by *Listeria monocytogenes,* a mainly respiratory Gram-positive species (Light et al., 2018; Rivera-Lugo et al., 2022). We found that *L. plantarum* NCIMB8826R requires *ndh2,* encoding a type-II NADH dehydrogenase, and conditionally requires *pplA,* predicted to encode a flavin-binding, membrane reductase, in the FLEET pathway to reduce extracellular ferric iron or a polarized anode (Tejedor-Sanz et al., 2022b; Tolar et al., 2022a).

Despite the benefits of EET for *L. plantarum* growth and intracellular energy, members of this species lack the capacity to synthesize either flavins or quinones (Brooijmans et al., 2009; Thakur et al., 2016). Whereas exogenous flavins are essential for *L. plantarum* growth under all conditions, EET reduction of insoluble ferrihydrite (iron(III) oxyhydroxide) and production of current in a bioelectrochemical reactor by *L. plantarum* NCIMB8826R are dependent on exogenous quinones (Tejedor-Sanz et al., 2022b). *L. plantarum* NCIMB8826R does contain several genes hypothesized to condense 1,4-dihydroxy-2-naphthoic acid (DHNA) with an isoprenoid polymer to form demethylmenaquinone (DMK), the membrane electron carrier used by *L. monocytogenes* (Light et al., 2018) and *Enterococcus faecalis* (Hederstedt et al., 2020) for EET. However, *L. plantarum* lacks the other genes required for a complete menaquinone biosynthesis pathway (Watthanasakphuban et al., 2021). Instead, exogenous DHNA, a quinone present in foods and made by other bacteria, was sufficient for stimulating *L. plantarum* EET metabolism (Tejedor-Sanz et al., 2022b; Tolar et al., 2022a).

To better understand the sources and impacts of quinones on *L. plantarum* and its EET metabolism, we investigated the quinone diversity and concentration ranges used by *L. plantarum* for EET, the effects of those compounds on *L. plantarum* growth, and the significance of quinones produced by other LAB for *L. plantarum* in food fermentations. Our results reveal how *L. plantarum* EET metabolism is adapted for exogenous environmental quinones and engages in quinone cross-feeding with other food fermentation bacteria. The findings have significance for understanding the ecological and physiological importance of EET in microorganisms that primarily rely on fermentation for energy conservation.

## Results

### 1. *L. plantarum* quinone selectivity for EET

We previously reported that EET reduction of insoluble ferrihydrite and production of current in a bioelectrochemical reactor by our model strain *L. plantarum* NCIMB8826R is dependent on the presence of exogenous DHNA in the assay medium (Tejedor-Sanz et al., 2022b). To determine whether this activity is specific to DHNA or if other ecologically-relevant quinones can be used, ferrihydrite reduction was measured after *L. plantarum* growth and subsequent incubation in mannitol (55 mM) as an energy source and different concentrations of either DHNA, ACNQ (2-amino-3-carboxy-1,4-naphthoquinone), menadione, hydroquinone, phylloquinone (phytomenadione, vitamin K1), 1,4-naphthoquinone, or the menaquinones MK-4 and MK-7. DHNA, ACNQ, and menadione are produced by bacteria (Altwiley et al., 2021; Mevers et al., 2019; Widhalm and Rhodes, 2016). The quinone precursor hydroquinone is found in both animal and fungal tissues (James et al., 2013; Joval et al., 1996). Phylloquinone and its precursor 1,4-naphthoquinone are contained in plant tissues (Widhalm and Rhodes, 2016). MK-4 and MK-7 are present in animal tissues and are also produced by bacteria (Walther and Chollet, 2017). In addition to DHNA, *L. plantarum* reduced ferrihydrite when ACNQ, 1,4-naphthoquinone, or menadione were included in the assay medium (**Figure 1**). ACNQ, a soluble analog of DHNA, resulted in the highest level of ferrihydrite reduction by *L. plantarum* with a maximum of 0.68 mM Fe^2+^ reduced and an IC_50_ of 4.70 μg/mL. *L. plantarum* also used 1,4-naphthoquinone (0.05 Fe^2+^ max, IC_50_ = 10.23 μg/mL) and menadione (0.05 Fe^2+^ max, IC_50_ = 29.79 μg/mL) to reduce iron, but less effectively than DHNA (0.40 Fe^2+^ max, IC_50_ = 223.4 μg/mL). In contrast, ferrihydrite was not reduced by *L. plantarum* in the presence of hydroquinone, phylloquinone, MK-4, or MK-7 up to the maximum concentration tested (100 μg/mL) (**Figure 1**). The high partitioning coefficients (LogP) of those quinones suggest that *L. plantarum* uses naphthoquinone-based, hydrophilic quinones for EET. These findings show that *L. plantarum* quinone requirements for EET are not limited to DHNA and that other environmental quinones are sufficient, providing a range of selectivity.

**Figure 1.**
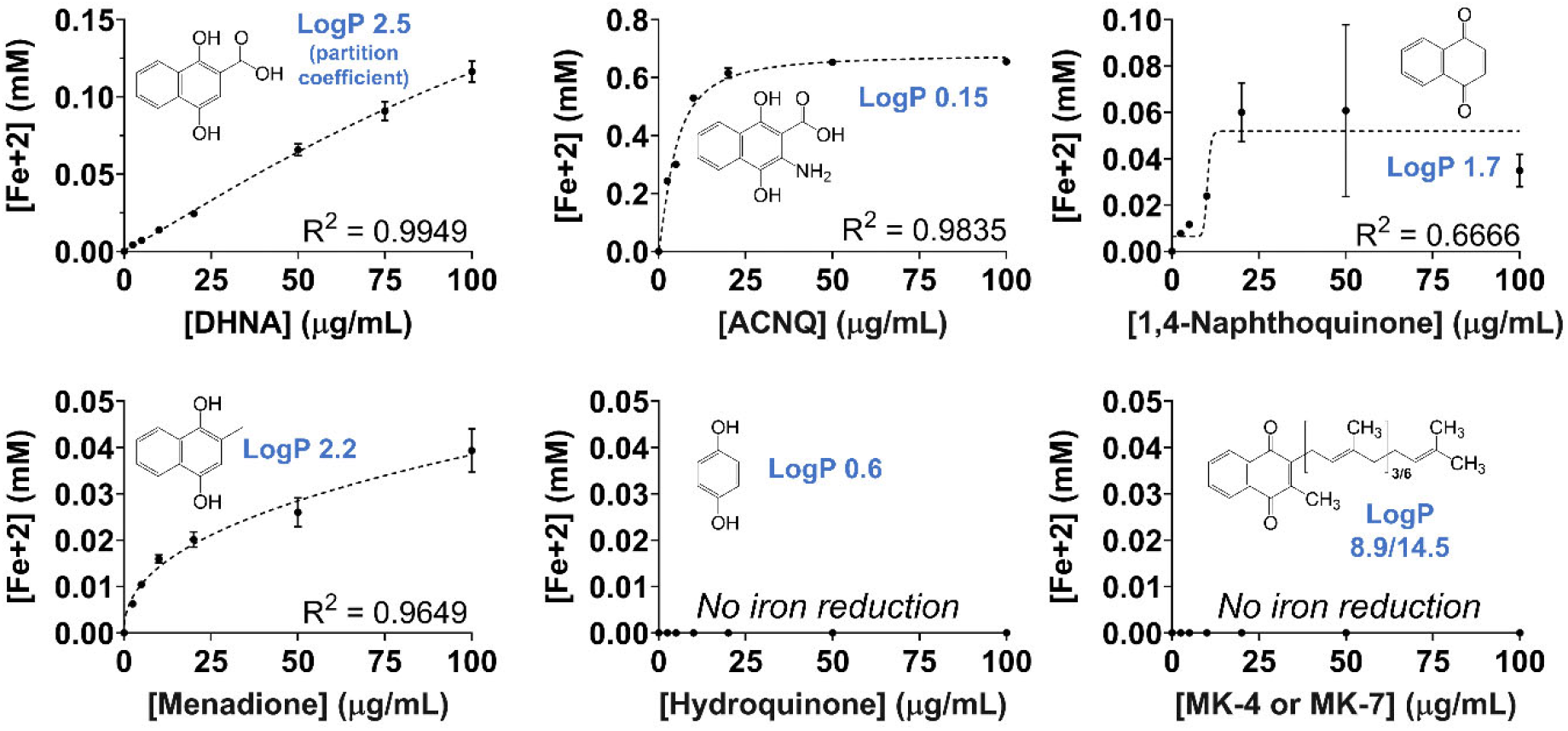
*L. plantarum* uses multiple quinone species for iron reduction. Ferrihydrite reduction assays were performed in PBS containing 55 mM mannitol and at the specified quinone concentrations. *L. plantarum* cells used in the assay were grown in mMRS without quinone supplementation. The R^2^ value corresponds to the sigmoidal dose-response curve fit to the data. The chemical structure of each quinone and its corresponding Log_P_ (partition coefficient) are provided. The avg ± stdev of three biological replicates is shown.

### Quinones induce oxidative stress inhibiting *L. plantarum* growth

If *L. plantarum* uses EET as a metabolic strategy in food fermentations, its growth should not be negatively impacted by antimicrobial activity associated with those compounds (Campos-Xolalpa et al., 2021). Contrary to this assertion, we observed that under the static, non-aerated growth conditions common for food fermentations, *L. plantarum* was inhibited by DHNA and growth rates significantly declined when DHNA was added to the laboratory culture medium (**Figure 2** and **Supplementary Figure S1**). In mMRS with 5 μg/mL DHNA, *L. plantarum* growth rates were reduced from 0.26 ± 0.01 h^-1^ (mMRS) to 0.22 ± 0.01 h^-1^ (p < 0.05) (**Supplementary Figure S1**). Growth rate decreased exponentially as a function of DHNA concentration and no growth was observed starting at 150 μg/mL DHNA. This effect was not due to activation of EET because growth of the *L. plantarum* Δ*ndh2* and Δ*pplA* mutants was also negatively affected when DHNA was included in mMRS (**Figure 2A**). Exponential growth inhibition also occurred during *L. plantarum* incubation in mMRS with the other EET-conducive quinones ACNQ, menadione, and 1,4-naphthoquinone (**Supplementary Figure S1**). There was no observable increase in *L. plantarum* cell numbers when either menadione or 1,4-naphthoquinone were added a concentration of 50 μg/mL. ACNQ was less toxic, and *L. plantarum* growth stopped at 100 μg/mL. Because ACNQ resulted in the greatest quantities of iron reduced (**Figure 1**), reductions in *L. plantarum* growth were not directly related to the use of the quinone as an electron shuttle for EET. These findings indicate that while *L. plantarum* can use different quinones as electron shuttles, exposure to these compounds could come at the cost of reducing ecological fitness.

**Figure 2.**
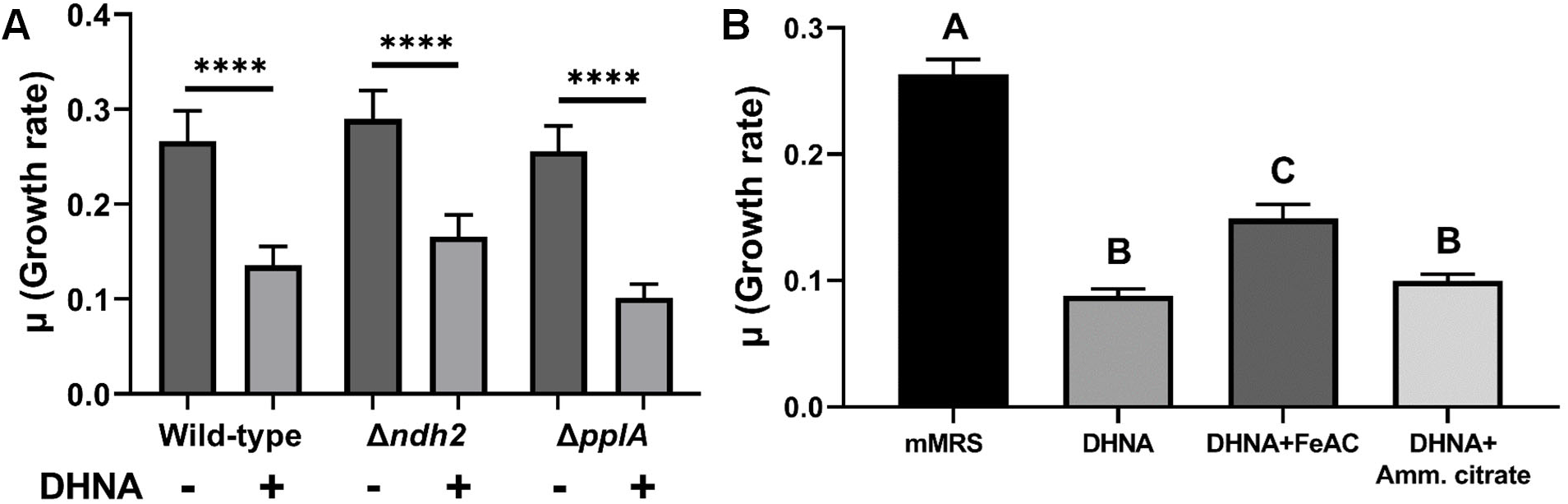
DHNA reduces *L. plantarum* growth and this is ameliorated by soluble ferric iron in the laboratory culture medium. **A)** Growth rate of wild-type *L. plantarum* NCIMB8826R and the Δ*ndh2* and *ΔpplA* mutants, MLES100 and MLES101, respectively, in mMRS with or without supplementation of 20 μg/mL DHNA. **B)** Growth rate of *L. plantarum* NCIMB8826R in mMRS with or without supplementation of 20 μg/mL DHNA and 1.25 mM ferric ammonium citrate (FeAC) or 1.25 mM ammonium citrate. Growth rates were quantified by measuring the change in OD_600nm_ per hour during exponential phase. Significant differences represented by asterisks (**** p ≤ 0.0001) and letters (p ≤ 0.05) were determined using one-way ANOVA with Tukey’s post-hoc test. The avg ± SEM of three biological replicates is shown.

To understand why growth was reduced in the presence of quinones, we measured the transcriptomic responses of exponential-phase *L. plantarum* cells in mMRS supplemented with 20 µg/ml DHNA. Compared to cells in mMRS, expression levels of approximately 30% of the genes in the *L. plantarum* genome (452 genes up-regulated and 473 genes down-regulated) were affected (**Table S1** and **Supplementary Table S1**). Pathway analysis of the transcriptionally induced genes showed that *L. plantarum* was undergoing oxidative stress (**Figure 3 and Supplementary Table S2**). Transcripts for the genes encoding glutathione (*gshR2*/*R3*/*R4*), methionine-S-oxide reductase (*msrA2* and *msrB*), and thioredoxin (*trxB*) reductases increased between 1.5-to 7-fold during incubation in DHNA. Likewise, genes for inactivating ROS including NADH oxidase (*nox5*), NADH peroxidase (*npr2*), catalase (*kat*), and pyruvate oxidase (*pox3*) were similarly upregulated. Expression levels of genes coding for amino acid and lipid metabolism were also significantly greater in the presence of DHNA. Transporters for sulfur-containing amino acids, as well as methionine and chorismate biosynthesis genes previously found to protect against oxidative stress (Bin et al., 2017a; Gerstle et al., 2012; Kim et al., 2020a; Zhang et al., 2012), were also induced (**Supplementary Table S2**). Induction of membrane biosynthetic pathways (acetyl-CoA carboxylase (*accA2*, *accB2*, *accC2*, and *accD2*) and fatty acid biosynthesis (*fabD*, *fabF*, *fabG*, *fabH2*, *fabI*, and *fabZ1*) was consistent with *E. coli* responses to lipid peroxidation (Janßen and Steinbüchel, 2014).

**Figure 3.**
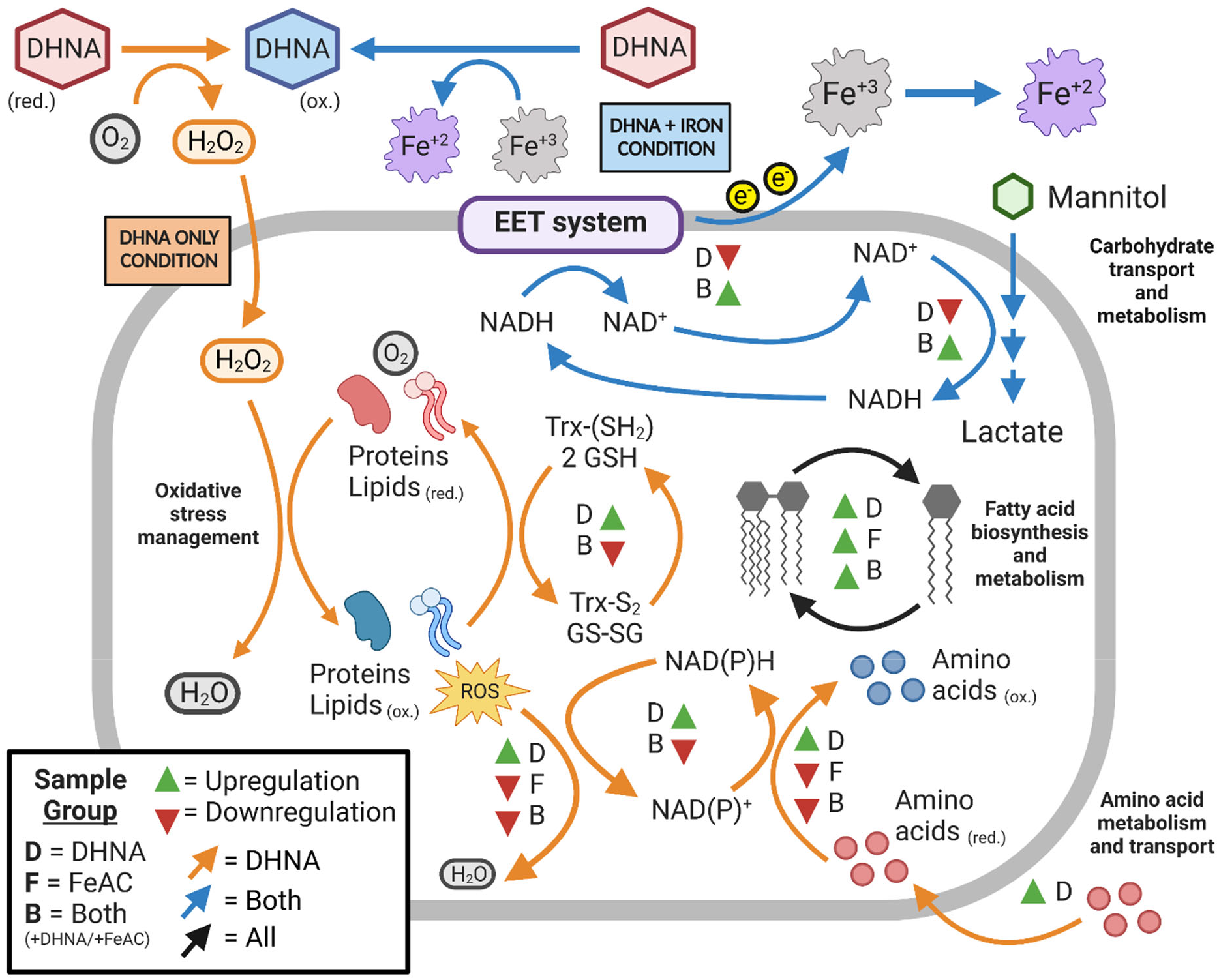
DHNA-caused induction of oxidative stress response in *L. plantarum* is alleviated by FeAC. Summary of. *plantarum* transcriptome changes in response to DHNA (D = DNHA), FeAC (F = FeAC), or DHNA+FeAC (B = Both) during exponential-phase growth in mMRS. Statistically significant changes in gene expression were determined using DEseq2 with a Log_2_ expression fold-change ≥ 0.5 and FDR-adjusted *p*-value ≤ 0.05.

Because the transcriptional responses of *L. plantarum* to DHNA signaled the activation of pathways responsive to intracellular oxidative stress, we measured for the presence of ROS in the mMRS medium. It was previously shown that hydroxyl groups of quinones react with O_2_ to result in the abiotic production of hydrogen peroxide (H_2_O_2_) (Wendlandt and Stahl, 2016). Although *L. plantarum* cultures were not aerated, in order to resemble the normal growth conditions of LAB in food environments, a strict anaerobic environment was also not intended. After 5 h incubation of *L. plantarum* in mMRS with DHNA, there was 29.92 mM ± 0.63 mM H_2_O_2_, a concentration approximately 15-fold higher than during *L. plantarum* growth in mMRS alone (**Supplementary Figure S2**). *L. plantarum* was not responsible for the elevated H_2_O_2_ concentrations because sterile mMRS+DHNA contained even higher levels of that compound (41.90 mM ± 0.34 mM H_2_O_2_). Thus, our data point to increased concentrations of H_2_O_2_ as the cause of *L. plantarum* growth rate reductions and increased oxidative stress responses in the presence of quinones.

### Electron acceptors alleviate DHNA induced stress

Our initial findings showed that including DHNA in laboratory culture medium together with soluble iron (ferric ammonium citrate, FeAC [1.25 mM]) as an exogenous electron acceptor greatly improved EET in the post-growth, ferrihydrite reduction assay (Tejedor-Sanz et al., 2022b). Consistent with FeAC increasing *L. plantarum* EET, the addition of this compound helped to alleviate some of the antimicrobial effects of DHNA and resulted in higher *L. plantarum* growth rates (0.15 ± 0.01 h^-1^) compared to when only DHNA was present (0.09 ± 0.01 h^-1^) (**Figure 2B**). The increase was due to the addition of iron and not citrate, because there was no improvement in growth when ammonium citrate was added instead (**Figure 2B**). Additionally, these effects of FeAC on *L. plantarum* were only observed when DHNA was also included in the mMRS. The growth rate of *L. plantarum* in the presence of FeAC was equivalent to the mMRS controls (0.12 ± 0.00 h^-1^) (**Supplementary Figure S3**). These data strongly suggest that iron reduces the oxidative stress generated by quinones in the presence of O_2_.

Consistent with higher growth rates of *L. plantarum* when FeAc was included together with DHNA in mMRS, the presence of FeAC resulted in lower H_2_O_2_ concentrations (8.34 ± 0.02 mM) (p ≤ 0.0001) (**Supplementary Figure S2**). Similarly, *L. plantarum* genes required for oxidative stress tolerance and redox-associated, amino acid metabolism were either unaffected or were downregulated during growth in mMRS with FeAC or mMRS with FeAC and DHNA (**Figure 3** and **Supplementary Table S2**). Transcriptional changes were not completely prevented with the addition of FeAC, however, because genes for lipid metabolism and cell membrane biosynthesis were upregulated. These data establish that *L. plantarum* growth is inhibited by exogenous quinones due to oxidative stress, and the detrimental effects of DHNA are largely diminished when a terminal electron acceptor is also present.

### Growth in DHNA and FeAC increases *L. plantarum* EET

Next, we aimed to understand the extent that EET is enhanced when *L. plantarum* is grown in the presence of DHNA and FeAC and why this occurs. Significantly more iron was reduced in the post-growth ferrihydrite assay when those compounds were included in the laboratory culture medium (**Figure 4A** and **Figure 4B**). The effect of DHNA and FeAC was observed over a broad range of concentrations (0.1 µg/ml to 200 µg/ml), and remarkably, the impact of adding it the growth medium was greatest when lower quantities of DHNA (< 20 µg/ml) were used (**Figure 4A**). Increased iron reduction was still observable when only 0.1 µg/ml DHNA was included (**Figure 4B**). The findings therefore indicate that exposure to FeAC and even low quantities of DHNA during growth induces physiologic changes to *L. plantarum* that dramatically enhance its capacity to perform EET.

**Figure 4.**
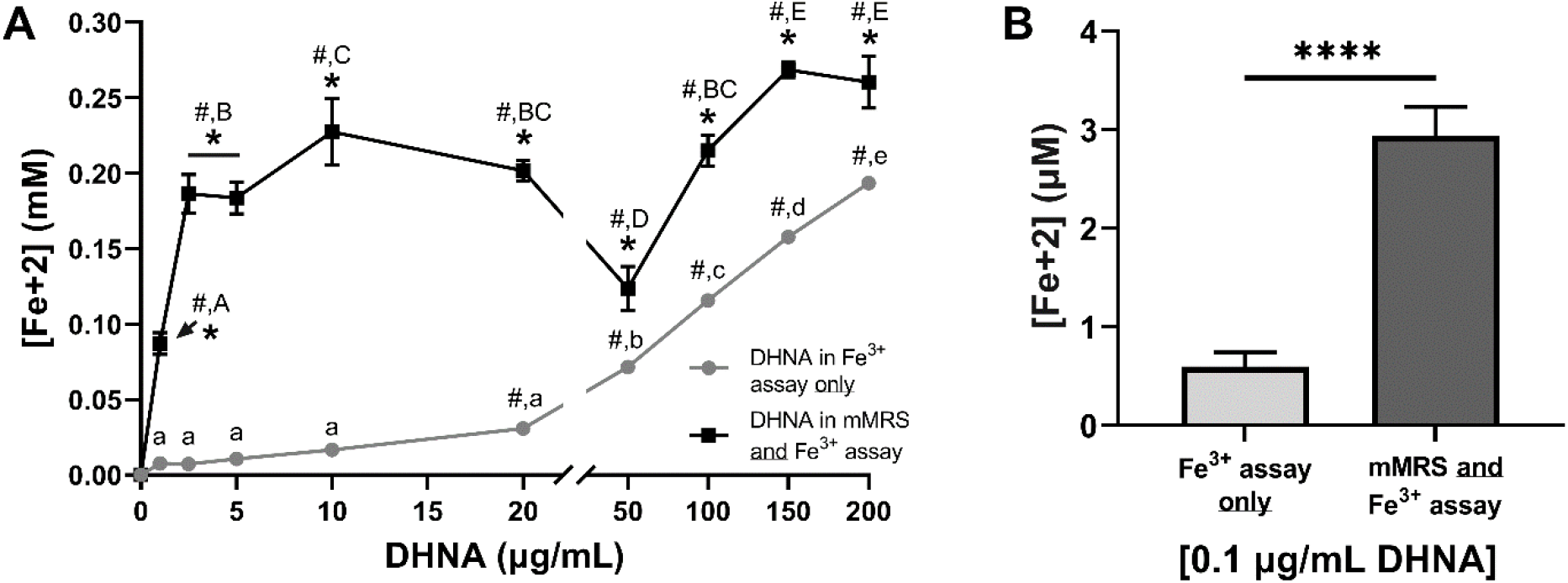
Growth in DHNA results in greater stimulation of *L. plantarum* EET. Reduction of Fe^3+^ (ferrihydrite) to Fe^2+^ by *L. plantarum* after growth in mMRS with 1.25 mM ferric ammonium citrate (FeAC) and DHNA. DHNA was supplemented in the growth medium and/or the ferrihydrite reduction assay reduction assay at the indicated concentrations (**A**) or at 0.1 µg/ml (**B**). Significant differences were determined by one-way ANOVA with Tukey’s post-hoc test (p ≤ 0.05). Letters denote when iron reduction was significantly different between DHNA concentrations within the same growth condition. Asterisks (*) denote when iron reduction was significantly greater (p ≤ 0.05) when the same concentration of DHNA was either included or excluded from the mMRS growth medium. Pound signs (#) denote when iron reduction was significantly greater (p ≤ 0.05) than when no DHNA was included in the ferrihydrite reduction assay. In panel B **** p ≤ 0.0001. The avg ± stdev of three biological replicates is shown.

One possible response that could improve the capacity of *L. plantarum* to perform EET is the upregulation of genes within the FLEET locus. Growth of *L. plantarum* mMRS with DHNA and FeAC was previously observed to result in the induction of the FLEET pathway genes *ndh2* and *pplA* (Tejedor-Sanz et al., 2022b) (**Supplementary Table S2**). However, other genes in the FLEET locus were not induced, potentially indicating that those genes are dispensable for EET. To this regard, an *L. plantarum* mutant lacking the FLEET locus genes encoding electron transfer proteins EetA and EetB, or the heptaprenyl diphosphate synthase DmkB, was still able to perform DHNA-dependent EET, albeit at a somewhat reduced level compared to the wild-type strain (Tolar et al., 2022a). Instead, genes needed for anaerobic respiration, namely those coding for nitrate reductase (*narGHJI*) and the molybdopterin cofactor (*moeA, moeB*), were induced in mMRS with DHNA and FeAC (**Supplementary Table S2**). While nitrate reductase is not required for the reduction of extracellular electron acceptors (outwards-EET) (Tejedor-Sanz et al., 2022b), the *narGHJI* operon was implicated in electron uptake from an electrode by *L. plantarum* in a biochemical reactor to result a metabolic shift towards ATP-generating pathways and prolonged survival after sugar exhaustion (Tejedor-Sanz et al., 2022a). Therefore, higher quantities of Ndh2, PplA, and nitrate reductase in *L. plantarum* after growth in mMRS containing DHNA and FeAC likely provided an increased EET capacity and an improved energetic state in the post-growth ferrihydrite reduction assay.

### *L. plantarum* converts DHNA to MK-6 and MK-7 but still relies on direct access to exogenous electron shuttles for EET

Transcriptomic analysis showed that genes required for menaquinone biosynthesis from DHNA, including *dmkA* (lp_1546), lp_1135, lp_1715, and *ubiE*, were not differentially expressed in either mMRS containing DHNA or both DHNA and FeAC (**Supplementary Table S2**). To determine whether DHNA could still be incorporated into *L. plantarum* cells irrespective of gene expression changes, intracellular quinones were measured by LCMS. After growth in mMRS with DHNA (with and without FeAC), menaquinone-6 (MK-6) and menaquinone-7 (MK-7) were the only quinones detected and present in an approximate ratio of 13 to 1 (8.67 x 10^6^ ± 6.53 x 10^5^ MK-6 ion counts and 6.53 x 10^5^ ± 3.56 x 10^5^ MK-7 ion counts, respectively). No quinones were detected when DHNA was omitted from the culture medium (data not shown). Because similar quantities of MK-6 and MK-7 were recovered from *L. plantarum* after growth in both EET-conducive (mMRS with DHNA and FeAC) and EET non-conducive conditions (mMRS with DHNA), it is expected that other factors besides the presence of intracellular quinones must be also involved in the heightened activity observed for *L. plantarum* collected from the EET-conducive growth conditions.

Another physiologic change that could increase EET metabolism is the secretion of quinones or other electron carriers into the extracellular environment. However, *L. plantarum* remained dependent on the addition of exogenous electron shuttles during the post-growth, ferrihydrite reduction assay (**Figure 5A**). These shuttles are not limited to DHNA because riboflavin was also used by *L. plantarum* to reduce ferrihydrite (**Figure 5A**). Moreover, in a bioelectrochemical reactor, *L. plantarum* generated current when DHNA was present, but this activity stopped when it was transferred to an electrochemical cell lacking an exogenous quinone source (**Figure 5B** and **Figure 5C**). The lack of a secreted soluble redox-active mediators was confirmed by cyclic voltammetry (**Figure 5D**), thus verifying that *L. plantarum* growth in the presence of DHNA and FeAC does not increase EET from the secretion of electron shuttles.

**Figure 5.**
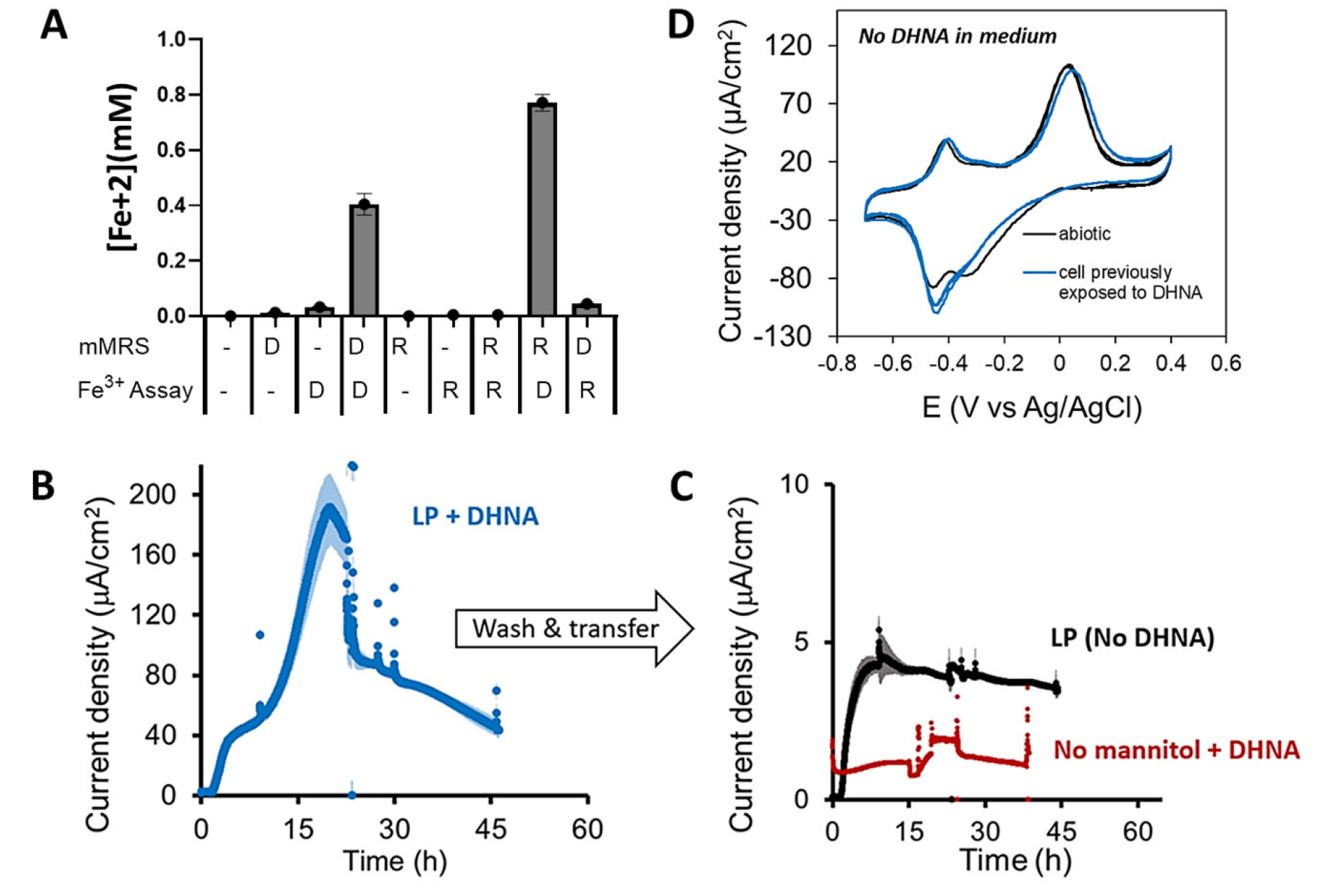
Access to exogenous electron shuttles is required for *L. plantarum* EET. **(A)** Ferrihydrite reduction measured after growth in mMRS with DHNA (D) or supplemental riboflavin (R) or inclusion of DHNA or riboflavin in the post-growth ferrihydrite (Fe^3+^) assay. (**B**). Current density production over time by *L. plantarum* in mCDM supplemented with 20 μg/mL DHNA. The anode was polarized at +0.2V Ag/AgCl. (**C**) Cells from (**B**) were washed and transferred to bioreactors containing mCDM with no DHNA or CDM with no mannitol or DHNA. (**D**) Cyclic voltammetry analysis of *L. plantarum* in mCDM with either prior growth in mCDM with 20 μg/mL DHNA or mCDM alone. The avg + stdev of three biological replicates is shown.

In summary, *L. plantarum* EET is inducible and more sensitive to low quantities exogenous electron shuttles when it is grown in the presence of DHNA and iron. *L. plantarum* converts DHNA to MK-6 and MK-7, but EET metabolism still requires additional exogenous electron shuttles such as quinones or flavins along with an environment containing electron acceptors like soluble iron (FeAC). Thus, although pathways used for DHNA conversion to MK-6 and MK-7 and signals needed for increased expression of *ndh2* and *pplA* are not yet known, growth in the presence of DHNA and FeAC resulted in significant increases in EET. Such increases may be stimulated by an increased energetic state, the availability of Ndh2 and PplA required for ferrihydrite reduction, reduced oxidative stress, and the presence of an intracellular quinone pool.

### *L. plantarum* uses quinones made by other LAB for EET

In food fermentations, *L. plantarum* is frequently found together with LAB known to synthesize quinones (Eom and Moon, 2015; Mevers et al., 2019; Morishita et al., 1999). To determine whether the secreted products from those LAB influence *L. plantarum* EET activity, we measured ferrihydrite reduction by *L. plantarum* during incubation in cell-free, spent-medium (CFS) collected after growth of the quinone-producer *L. lactis* TIL46 and a quinone-deficient, *menC* deletion mutant of that strain, TIL999 (Tachon et al., 2010) (**Supplementary Figure S4A**). Measurements of *L. lactis* intracellular quinones following growth in medium lacking exogenous quinone sources (GM17) showed that TIL46 contained MK-7, MK-8, and MK-9 in a ratio of 1:7:9. Menaquinone levels were significantly reduced (approximately 16-fold) in TIL999 (**Table 2**). Consistent with the quinone biosynthetic capacity of *L. lactis* TIL46, significantly greater ferrihydrite reduction was observed for *L. plantarum* incubated in *L. lactis* TIL46 CFS than TIL999 CFS (**Figure 6A**). The lack of activity when *L. plantarum* Δ*ndh2* was incubated with the CFS from either *L. lactis* strain confirmed that the increase in iron reduction was dependent on an intact *L. plantarum* FLEET pathway (**Figure 6A**). To assess whether *L. plantarum* EET stimulation is limited to *L. lactis* CFS, we also examined CFS from *Leuconostoc mesenteroides* ATCC8293. *L. mesenteroides* cells contained MK-10 and MK-11, present in a ratio of 1:3 (**Table 2**). CFS from *L. mesenteroides* ATCC8293 also enabled *L. plantarum* EET (**Figure 6B**). Like found for *L. lactis* TIL46, this activity was dependent on a functional *L. plantarum ndh2* (**Figure 6B**).

**Table 1.** Strains and plasmids used in this study.

| Species | Strain | Description | Reference |
| --- | --- | --- | --- |
| *L. plantarum* | NCIMB8826R | Human saliva | (Dandekar, 2019) |
| *L. plantarum* | MLES100 | Deletion mutant of NCIMB8826 lacking *ndh2* | (Tejedor-Sanz et al., 2022b) |
| *L. plantarum* | MLES101 | Deletion mutant of NCIMB8826 lacking *pplA* | (Tejedor-Sanz et al., 2022b) |
| *L. lactis subsp. cremoris* | TIL46 | Strain NCDO763 cured of its 2-kb plasmid | (Chambellon and Yvon, 2003) |
| *L. lactis subsp. cremoris* | TIL999 | Deletion mutant of TIL46 lacking *menC* | (Tachon et al., 2009) |
| *L. mesenteroides subsp. mesenteroides* | ATCC8293 | Fermented olives | (Makarova et al., 2006) |

**Table 2.** Cellular menaquinones in lactic acid bacteria.

| Strain | Menaquinone production normalized to cell pellet |  |  |  |  |  |
| --- | --- | --- | --- | --- | --- | --- |
|  | MK6 | MK7 | MK8 | MK9 | MK10 | MK11 |
| *Lactiplantibacillus plantarum*<br>NCIMB8826R | 107.28 ± 23.07 | 11.29 ± 7.51 | ND | ND | ND | ND |
| *Lactococcus lactis* TIL46 | 151.09 ± 32.87 | 471.12 ± 172.65 | 3265.06 ± 1244.88 | 4297.39 ± 1904.09 | 261.46 ± 146.37 | ND |
| *Lactococcus lactis* TIL999 | 8.92 ± 5.15 | ND | 8.90 ± 2.85 | 109.60 ± 118.46 | ND | ND |
| *Leuconostoc mesenteroides*<br>ATCC8293 | ND | ND | 0.08 ± 0.15 | 4.28 ± 2.38 | 124.94 ± 34.87 | 312.42 ± 97.64 |

Mean peak area per gram cell pellet ± stdev (n = 3) is provided. ND = None detected

**Figure 6.**
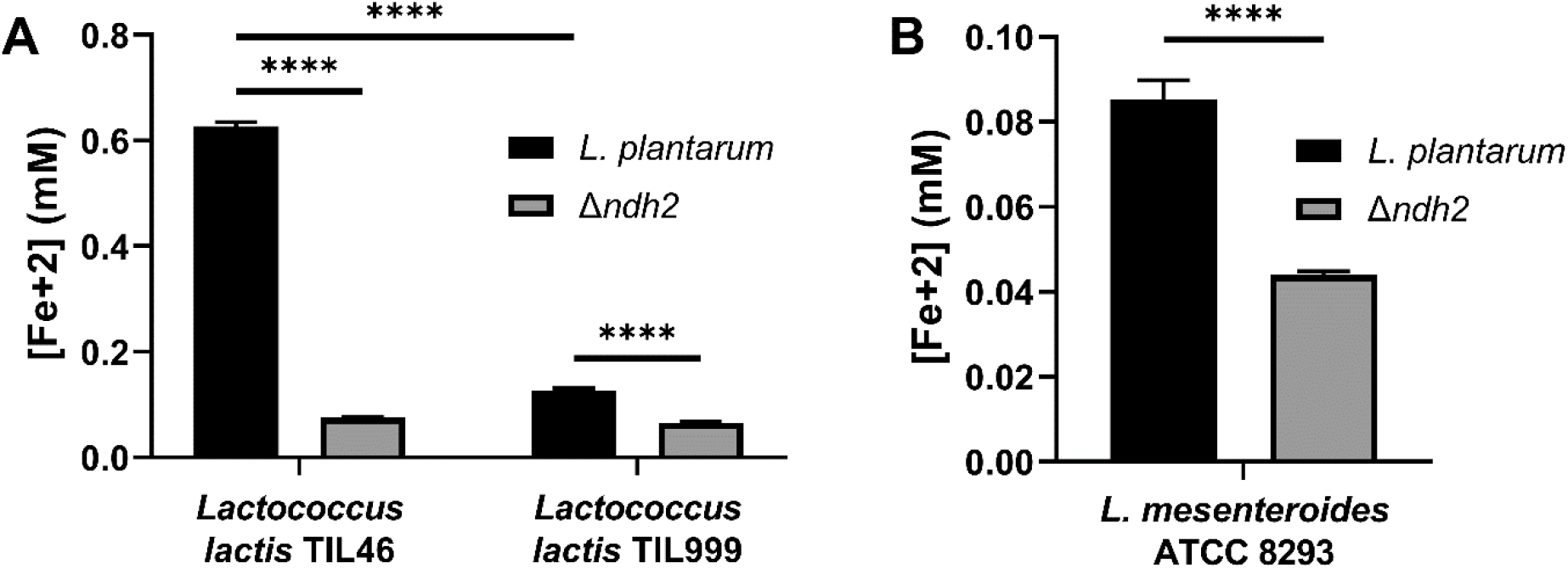
*L. plantarum* EET is activated by secreted compounds made by *L. lactis* and *Leuconostoc* in an EET-dependent manner. Ferrihydrite reduction by wild-type *L. plantarum* and the *ndh2* deletion mutant MLES100 in cell-free supernatant (CFS) from (**A**) wild-type *L. lactis* TIL46 and the *menC* deletion mutant TIL999 or (**B**) *L. mesenteroides* ATCC8293. Significant differences between groups (*** p ≤ 0.001; **** p ≤ 0.0001) were determined by one-way ANOVA with Tukey’s post-hoc test. The avg ± stdev of three biological replicates is shown.

### Quinone cross-feeding by *L. plantarum* increases EET and environmental acidification

Because the CFS from quinone-producing LAB was sufficient for *L. plantarum* EET, we hypothesized that quinone cross-feeding in co-culture would also simulate *L. plantarum* iron reduction *in situ*. *L. plantarum* was incubated in equal proportions with either *L. lactis* TIL46, *L. lactis* TIL999, or *L. mesenteroides* ATCC8293 in a chemically defined medium (gCDM) and 1.25 mM FeAC. The addition of FeAC into that medium also provided a terminal electron acceptor that we could use to measure EET activity *in situ* via colorimetric analysis (**Supplementary Figure S4B**). Although there was spontaneous reduction of FeAC in acidified gCDM (**Supplementary Figure S5**), this background level of iron reduction was minor and did not impact the significant increases in iron reduction when *L. plantarum* was present.

After incubation of *L. plantarum* and *L. lactis* TIL46 in co-culture for 6 h, significantly more iron (0.149 ± 0.19 mM Fe^2+^) was reduced than when either *L. lactis* TIL46 (0.056 ± 0.01 mM Fe^2+^) or *L. plantarum* was grown separately (**Figure 7A**). No iron reduction was found following TIL999 growth or the co-culture of *L. plantarum* and TIL999 (**Figure 7A**). By 24 h, *L. lactis* TIL46 and the co-culture of *L. plantarum* and *L. lactis* TIL46 reduced similar amounts of iron (0.43 ± 0.02 mM Fe^2+^) (**Figure 7A**). These quantities were significantly greater than found for either *L. plantarum* or TIL999 grown alone (0.075 ± 0.02 mM Fe^2+^). Although FeAC was reduced in the co-culture of *L. plantarum* and TIL999 (0.197 ± 0.00 mM Fe^2+^) the levels were significantly lower compared to cultures containing the *L. lactis* wild-type strain. We previously reported that the genomes of *L. lactis* strains lack *pplA* and that this gene is required for ferrihydrite reduction in the post-growth assay (Tejedor-Sanz et al., 2022b). However, the findings here show that *L. lactis* is able to reduce FeAC *in situ* and this activity is partially dependent on its quinone biosynthetic capacity. For co-culture of *L. plantarum* and *L. mesenteroides*, FeAC reduction was found after incubation for 6 h (0.01 ± 0.00 mM Fe^2+^), whereas no iron was reduced by either strain grown separately (**Figure 8A**). After 24 h, this value increased in the co-culture (0.19 ± 0.00 mM Fe^2+^) and remained significantly higher than the *L. plantarum* (0.13 ± 0.00 mM Fe^2+^) or *L. mesenteroides* (0.11 ± 0.01 mM Fe^2+^) cultures (**Figure 8A**). These results show that environmental quinones secreted by other LAB can stimulate *L. plantarum* EET metabolism, and that full quinone biosynthetic capabilities are necessary for robust iron reduction by *L. lactis*.

**Figure 7.**
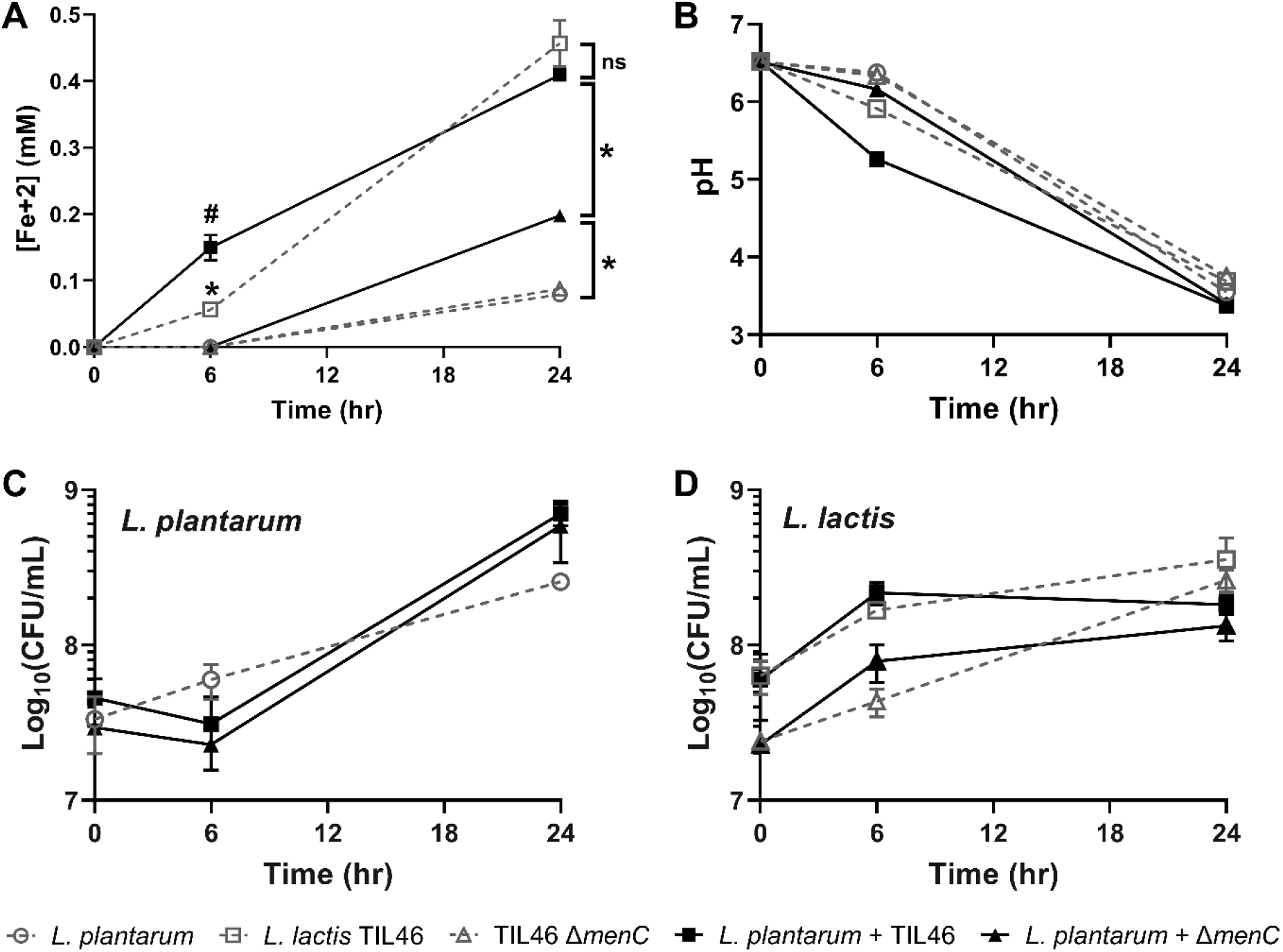
Co-culturing *L. plantarum* with quinone-producing *L. lactis* increases EET, acidification and *L. plantarum* growth. (**A**) Ferrihydrite reduction by *L. plantarum* in gCDM with 1.25 mM FeAC during growth alone or in co-culture with *L. lactis* TIL49 or TIL999. Change in (**B**) pH and (**C**) *L. plantarum* and (**D**) *L. lactis* abundance over time when grown separately or combined in co-culture. Strains were grown in gCDM supplemented with 1.25 mM FeAC. Significant differences in strain abundance were determined by two-way ANOVA with Tukey’s post-hoc test (*,^#^ p ≤ 0.05). The avg ± stdev of three biological replicates is shown.

**Figure 8.**
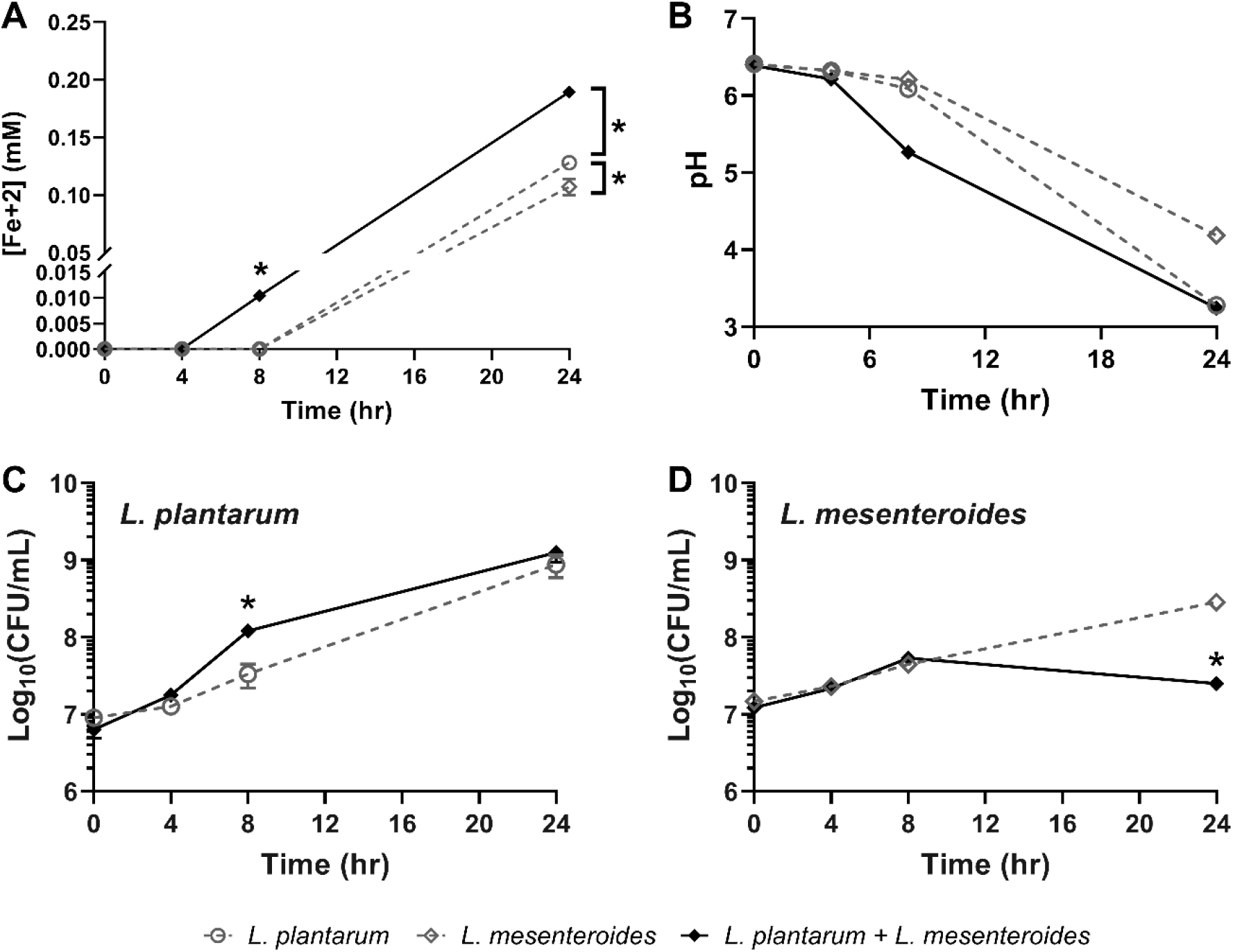
Co-culturing *L. plantarum* with quinone-producing *L. mesenteroides* increases EET, acidification, and *L. plantarum* growth. (**A**) Ferrihydrite reduction by *L. plantarum* in gCDM with 1.25 mM FeAC during growth alone or in co-culture with *L. mesenteroides* ATCC8293. Change in (**B**) pH and (**C**) *L. plantarum* and (**D**) *L. mesenteroides* abundance over time when grown separately or combined in co-culture. Strains were grown in gCDM supplemented with 1.25 mM FeAC. Significant differences in strain abundance were determined by two-way ANOVA with Tukey’s post-hoc test (* p ≤ 0.05) The avg ± stdev of three biological replicates is shown.

*L. plantarum* EET is associated with a shorter lag phase, increased viable cell counts, and accelerated environmental acidification (Tejedor-Sanz et al., 2022b). Because incubation of *L. plantarum* with quinone-producing LAB promoted EET metabolism, we also quantified environmental acidification and strain growth in co-culture. Within 6 h of incubation, medium acidification was greater for the *L. plantarum* and *L. lactis* TIL46 co-culture (pH = 5.26 ± 0.03) than the *L. plantarum* and TIL999 co-culture (pH = 6.16 ± 0.03) as well as for *L. plantarum* (pH = 6.4 ± 0.01), TIL46 (pH = 5.91 ± 0.05), and TIL999 (pH = 6.34 ± 0.02) grown separately (**Figure 7B**). Likewise, co-cultures of *L. plantarum* and *L. mesenteroides* ATCC8293 led to greater acidification of the medium (pH = 5.27 ± 0.05) within 8 h, compared to when the strains were grown separately (pH = 6.10 ± 0.01 and pH = 6.21 ± 0.01 for *L. plantarum* and *L. mesenteroides*, respectively) (**Figure 8B**). These differences were no longer observed after 24 h incubation (**Figure 7B** and **Figure 8B**), consistent with previous observations that *L. plantarum* EET is active during the early exponential phase of growth (Tejedor-Sanz et al., 2022b). Hence, the data show that quinone-cross feeding from LAB to *L. plantarum* results in accelerated environmental acidification.

Besides the increased acidification rate, strain growth was altered in the *L. lactis* and *L. mesenteroides* co-cultures. When *L. lactis* was present, *L. plantarum* cell numbers were 2-fold higher after incubation for 24 h (**Figure 7C**). However, this increase was not significant nor was it dependent on menaquinone biosynthetic capacity because both TIL46 and TIL999 resulted in similar increases in *L. plantarum* cell quantities (**Figure 7C**). Growth of *L. lactis* was not affected by *L. plantarum* (**Figure 7D**). Conversely, *L. plantarum* cell numbers increased 4-fold upon 8 h incubation with *L. mesenteroides* compared to when it was grown separately (p = 0.03) (**Figure 8C**). Growth of *L. mesenteroides* was negatively affected by co-culture and 10-fold lower cell numbers were found after 24 h incubation with *L. plantarum* (**Figure 8D**). Hence, *L. plantarum* exhibits antagonistic behavior against *L. mesenteroides* in a manner that is consistent with its EET-mediated metabolism.

## Discussion

Quinones are abundant in fermented foods, human and animal microbiomes, as well as other environments, where they may be responsible for important ecological and physiological processes (Franza and Gaudu, 2022; Karl et al., 2017; Walther and Chollet, 2017). We found that *L. plantarum,* a quinone auxotroph, can use multiple quinone species, including quinones made by other related bacteria, for EET. When an electron acceptor is present in food fermentation-relevant conditions, *L plantarum* partially overcomes the growth inhibiting effects of DHNA. *L plantarum* incorporates quinones into its intracellular contents when DHNA is present, but still requires direct access to those compounds or other electron shuttles for EET. Increased, initial environmental acidification during co-culture with *L. lactis* and *L. mesenteroides* and changes to *L. plantarum* and *L. mesenteroides* growth show the ecological significance of EET for these mainly fermentative bacteria.

### L. plantarum uses hydrophilic exogenous quinones for EET

*L. plantarum* uses hydrophilic, naphthoquinone-based quinones with electron-withdrawing substituents such as menadione, ACNQ, and DHNA For EET. Many extracellular electron shuttles are small, hydrophilic molecules (Lin et al., 2021), and the presence of electron-withdrawing groups on quinones can stabilize radicals in the aromatic ring (Mirkhalaf et al., 2004). EET activity increased logarithmically with quinone concentrations. For DHNA, the range of quinone use is comparable to concentrations found in makgeolli, a fermented Korean rice wine (0.089 to 0.44 <u>μg/mL)</u> (Eom et al., 2012) and produced by dairy-associated bacteria including *Propionibacterium freudenreichii* (36.75 μg/mL) (Furuichi et al., 2006) and *Lacticaseibacillus casei* (0.37 μg/mL). Less is known about the quantities of menadione and ACNQ in *L. plantarum*-associated environments. However, *L. lactis* was previously shown to synthesize ACNQ (Freguia et al., 2009; Mevers et al., 2019) and a *S. oneidensis* was found to produce up to 0.33 μg/mL (Mevers et al., 2019). Consistent with these results, 138 μg/mL menadione and 0.02 μg/mL ACNQ stimulated EET activity in the LAB *E. faecalis* (Pankratova et al., 2018) and *L. lactis* (Freguia et al., 2009), respectively. These data are also supported by our previous findings showing that ferrihydrite reduction occurred when 20 μg/mL DHNA was provided to several other LAB also known to be quinone auxotrophs, namely *Lactiplantibacillus pentosus*, *L. casei*, and *Lacticaseibacillus rhamnosus* (Pedersen et al., 2012; Tejedor-Sanz et al., 2022b).

*L. plantarum* EET was not observed when long-chain, hydrophobic menaquinones or phylloquinone were provided. This differs from results showing that *S. oneidensis* can use MK-4 as well as ACNQ, DHNA, and menadione for EET at ng/uL concentrations (Mevers et al., 2019). Similarly, although the CFS from *L. lactis* TIL46 and *L. mesenteroides* supported *L. plantarum* EET in a quinone-dependent manner, the intracellular contents of those LAB also contained the long-chain menaquinones MK-7 through MK-11. While it is possible that some of those MKs could be used by *L. plantarum* for EET, it is also possible that MK intermediates or derivatives were released from *L. lactis* or *L. mesenteroides* during growth, but not measured by LCMS targeting intercellular quinone contents. To this regard, extracellular ACNQ was found in spent media from *L. lactis* (Mevers et al., 2019). Other LAB genera including *Leuconostoc, Weissella,* and *Enterococcus* are also capable of synthesizing DHNA (Eom and Moon, 2015) or menaquinones (Pedersen et al., 2012).

### Quinones induce oxidative stress inhibiting L. plantarum growth, but this is alleviated by the addition of soluble iron

DHNA, ACNQ, menadione, and 1,4-naphthoquinone inhibited *L. plantarum* growth rates in a concentration-dependent manner. While evidence of quinone-induced growth inhibition on LAB is limited, DHNA, menadione, and 1,4-naphthoquinone were previously shown to possess antimicrobial activity against pathogens such as *Helicobacter pylori*, *Staphylococcus aureus*, and *Bacillus anthracis* (Nagata et al., 2010; Schlievert et al., 2013). *L. plantarum* growth inhibition was likely due the oxidation of O_2_ by these quinones resulting in H_2_O_2_ formation (Wendlandt and Stahl, 2016). Although *L. plantarum* and other lactobacilli possess genes for heme-dependent catalases, all LAB including *L. plantarum* are auxotrophic for heme (Zotta et al., 2017). Furthermore, *L. plantarum* lacks a functional manganese pseudo-catalase (Watanabe et al., 2012). *L. plantarum* instead relies on intracellular H_2_O_2_-detoxification enzymes like NADH peroxidase, glutathione reductase, and thioredoxin reductase to prevent oxidative stress (Zotta et al., 2017). These genes were upregulated in *L. plantarum* NCIMB8826R grown in mMRS with DHNA, and a similar oxidative stress response was observed for *L. plantaruim* CAUH2 upon exposure to 5 mM H_2_O_2_ (Zhai et al., 2020). The upregulation of sulfur-containing amino acid transporters during growth in mMRS with DHNA is also consistent with an oxidative stress response. Both methionine and cysteine are radical-scavenging amino acids (Kim et al., 2020b), cysteine is incorporated into glutathione, (Bin et al., 2017b), and methionine is incorporated in S-adenosylmethionine for cystathionine production (Sperandio et al., 2005). Moreover, the induction of *L. plantarum* genes required for chorismate biosynthesis, a precursor compound to the membrane-associated antioxidant ubiquinol (Gerstle et al., 2012), further indicates a protective response to oxidative conditions present in mMRS with DHNA.

The growth rate of *L. plantarum* was significantly improved and extracellular H_2_O_2_ was reduced when FeAC was included with DHNA in the laboratory culture medium. The majority of oxidative stress responses were downregulated when FeAC was present. The presence of FeAC alone had minor effects on *L. plantarum* growth and gene expression. Even though *L plantarum* can accumulate approximately 100-times higher quantities of iron during growth in FeAC (Tejedor-Sanz et al., 2022b), expression levels of *sufB* and *sufC* encoding iron-sulfur cluster assembly proteins used for detoxifying ROS in other LAB species (Papadimitriou et al., 2016), were down-regulated in the FeAC-containing medium. This result is surprising because ferric iron was shown to catalyze ROS production in the presence of quinones (Lyngsie et al., 2018). In mMRS, the complexing of iron with DHNA may make both compounds less available to react with molecular oxygen for hydrogen peroxide production (Tuchagues and Hendrickson, 1983). Thus, the addition of the terminal electron acceptor FeAC provided a means to reduce oxidative stress by lowering ROS synthesis.

### L. plantarum cells acquire quinones but still need access to environmental electron shuttles for EET

We found that although *L. plantarum* is a quinone auxotroph, it was still able to reduce extracellular ferrihydrite when incubated with DHNA and other hydrophilic quinones in the post-growth assay for several hours, and without prior inclusion of DHNA in the culture medium (**Figure 1** and **Figure 5A**). Because membrane-bound quinones are thus far known to be essential for EET in many bacteria (Glasser et al., 2017), it is possible that some quinone uptake occurred during that time. However, 5-fold higher levels of ferrihydrite were reduced when *L. plantarum* was first grown in laboratory culture medium containing DHNA and FeAC. That medium resulted in the induction of *ndh2* and *pplA* and the capacity of *L. plantarum* to use either DHNA or riboflavin as electron shuttles to reduce ferrihydrite. This increased EET capacity is likely due at least in part to the uptake and conversion of the exogenous quinone into its cellular contents. Because *L. plantarum* is able to respire using MK-4 for electron transport (Brooijmans et al., 2009), these bacteria are apparently able to acquire different quinone species for intracellular electron transfer. Similarly, *L. monocytogenes* mutants unable to make either menaquinone or demethylmenaquinone (DMK) lost the capacity for either aerobic respiration or EET metabolism, respectively (Light et al., 2018).

*L. plantarum* intracellular quinones, encompassing the long chain menaquinones MK-6 and MK-7, cannot be used by *L. plantarum* as extracellular electron shuttles for EET. These quinones also do not appear to be subsequently secreted because no soluble mediator was found in the culture media according to cyclic voltammetry or upon transfer to *L. plantarum* into other media. This is unlike other bacteria such as *Enterobacter cloacae*, an organism that self-secretes hydroquinones as an EET electron shuttle (You et al., 2019). Although the pathway for quinone incorporation into *L. plantarum* cells remains to be identified, it is notable that *L. plantarum* genes annotated for quinone metabolism were not induced when DHNA was included in the mMRS medium. While it is possible that the genes for menaquinone biosynthesis are constitutively expressed, it was also recently found that *L. plantarum* mutants lacking *dmkA* or *dmkB,* encoding a 1,4-dihydroxy-2-naphthoate polyprenyltransferase and heptaprenyl diphosphate synthase, respectively, were still able to perform EET in the presence of exogenous quinone (Tolar et al., 2022b). Therefore, it is expected that other, yet to be identified, pathways must be used by *L. plantarum* for quinone uptake and modification.

### Quinone cross-feeding to L. plantarum has important ecological implications

Incubation of *L. plantarum* with spent media (CFS) from either *L. lactis* TIL46 or *L. mesenteroides* ATCC8293 showed that *L. plantarum* EET was stimulated by the presence of the other strains. Reduction of FeAC increased 1.6-to 4-fold when CFS from *L. mesenteroides* and *L. lactis* TIL46 were provided. Although both *L. lactis* and *L. mesenteroides* are also capable of producing endogenous flavins (Thakur et al., 2016) that can be utilized as environmental electron shuttles (Tolar et al., 2022a), the capacity of *L. lactis* to synthesize quinones was required for *L. plantarum* EET stimulation. Notably, *L. lactis,* and to a lesser extent *L. mesenteroides*, were able to reduce FeAC on their own. Neither *L. lactis* nor *L. mesenteroides* possess a complete FLEET pathway (Tejedor-Sanz et al., 2022b). In addition, although *L. lactis* could generate an electric current, it was unable to reduce ferrihydrite in the post-growth assay (Tejedor-Sanz et al., 2022b). This reinforces that there are multiple routes for extracellular reduction in LAB besides using FLEET. Moreover, the observation that modest EET activity was observed with TIL999 shows there are other electron shuttles generated (e.g. flavins) by *L. lactis* that can be used by *L. plantarum* for EET, albeit with reduced efficiency.

Only co-cultures containing *L. plantarum* and *L. lactis* TIL46 or *L. plantarum* and *L. mesenteroides* resulted in an accelerated environmental acidification. Both co-cultures reached similar pH levels at during early exponential growth (between 6 to 8 h incubation). The pH change was correlated with quinone biosynthesis because the pH reductions were not observed upon *L. plantarum* incubation with the *menC* deletion mutant TIL999. These results are consistent with the observed use of EET by *L. plantarum* during growth in kale juice (Tejedor-Sanz et al., 2022b). In EET-conducive conditions with a polarized anode, there was a greater reduction in extracellular pH resulting from an increase in metabolic flux and lactic acid production (Tejedor-Sanz et al., 2022b). Like our previous results, this rapid acidification profile from EET was transient and after 24 h growth similar pH values were reached for *L. plantarum* grown separately or together with the other LAB strains.

The co-culture with *L. mesenteroides* also resulted in an initial increase in *L. plantarum* growth rate, inferred by the higher cell numbers within the first 8 h of incubation. Similarly, transient increases in *L. plantarum* cell numbers were observed in the kale juice medium under EET-conducive relative to non-conductive conditions (Tejedor-Sanz et al., 2022b). Although the lack of change in *L. plantarum* cell numbers during incubation with either the *L. lactis* TIL46 or TIL999 indicates that quinone-independent factors are likely influencing competition between *L. plantarum* and *L. lactis,* other studies have shown positive outcomes for quinone-cross feeding. Group B *Streptococcus* (GBS) shifted from fermentation to respiration and exhibited improved growth and survival when grown together with quinone-producing strains of *L. lactis* (Rezaïki et al., 2008) or *E. faecalis* (Franza and Gaudu, 2022). DHNA, MK-4, and menadione were found to stimulate the growth of strictly anaerobic bacteria isolated from the human intestine (Fenn et al., 2017). Similarly, DHNA produced by *Propionibacterium* increased growth (ΔA_580nm_ after 16 h growth) of *Bifidobacterium*, a bacterial genus that uses fermentation energy conservation metabolism (Isawa et al., 2002). Therefore, these findings suggest that quinone-cross feeding may be pervasive in food and intestinal habitats and that quinones are nutrients important for determining population dynamics in ecosystems wherein fermentation metabolism is prevalent.

## Conclusions

In summary, our results show there is a major metabolic and ecological impact of EET induced by exogenous quinones. These findings are important for the understanding microbial habitats such as fermented foods and animal digestive tracts containing microorganisms using fermentation for energy conservation. Many LAB are particularly well-adapted for those habitats, and because those sites also tend to be nutrient-rich, LAB have undergone reductive genome evolution and correspondingly rely on exogenous nutrients (amino acids, nucleosides, etc.) for growth (Makarova et al., 2006). The capacity of *L. plantarum* to use environmental quinones for EET is consistent with its adaptation to those nutrient-rich environments and may afford an increased competitive fitness compared to microorganisms that make those compounds de novo. The elucidation of the intracellular and extracellular quinone species needed for EET and the signals used by *L. plantarum* to induce the EET pathway will be valuable to apply these pathways to improve food fermentations and other biotechnological applications.

## Materials and Methods

### Strains and culture conditions

All strains and plasmids used in this study are listed in **Table S1**. Standard laboratory culture medium was used for routine growth of bacteria as follows: *Lactiplantibacillus plantarum* and *Leuconostoc* spp., MRS (BD, Franklin Lakes, NJ, USA), *Lactococcus* lactis, M17 (BD) with 2% w/v glucose (GM17), and *Escherichia coli*, LB (Teknova, Hollister, CA, USA). *L. plantarum* FLEET deletion mutants of *ndh2* or *pplA* were constructed through double-crossover homologous recombination as previously described (Tejedor-Sanz et al., 2022b). Strains were also grown at 30 or 37 °C when indicated. In place of standard laboratory culture medium, strains were grown (when indicated) in modified MRS (De Man et al., 1960) with no beef extract and with either 110 mM glucose [gMRS] or 110 mM mannitol [mMRS], or a chemically defined minimal medium (Tejedor-Sanz et al., 2022b) with 125 mM glucose [gCDM] or 125 mM mannitol [mCDM].

### L. plantarum growth rate experiments

After overnight growth in MRS at 37 °C, *L. plantarum* cells were collected via centrifugation (10,000 g / 3 min) and washed twice in phosphate-buffered saline (PBS) at pH 7.2 (http://cshprotocols.cshlp.org). Cells were resuspended in PBS at an optical density at 600 nm (OD_600nm_) of 1 before inoculation into 96-well culture plates (Thermo Fisher Scientific, Waltham, MA) at a final OD_600nm_ of 0.01 in mMRS. When indicated, the mMRS was supplemented with ferric ammonium citrate (VWR, Radnor, PA, USA) (1.25 mM) or ammonium citrate tribasic (Alfa Aesar, Haverhill, MA, USA) (1.25 mM). The following quinones were also supplemented; 1,4-benzenediol (hydroquinone) (Sigma-Aldrich, St. Louis, MO, USA), 1,4-dihydroxy-2-naphthoic acid (DHNA) (Alfa Aesar), 1,4-naphthoquinone (TCI, Tokyo, Japan), 2-methyl-1,4-naphthoquinone (menadione) (Sigma-Aldrich), Vitamin K1 (phylloquinone) (TCI), or Vitamin K2 in the form of menaquinone-4 (Sigma-Aldrich) or menaquinone-7 (Sigma-Aldrich). ACNQ was provided by E. Mevers and was prepared as previously reported (Mevers et al., 2019). Quinones were supplemented at 2.5, 5, 10, 20, 50, 100, 150, or 200 μg/mL where indicated. The OD_600nm_ was measured over 48 h in a Synergy 2 microplate reader (Biotek, Winooski, VT) set to 37 °C with no aeration.

In experiments where cell-free supernatant (CFS) was used in place of quinone supplementation, overnight cultures of *L. lactis* (GM17) or *Leuconostoc* spp. (gMRS) were grown for 18 h at 30 °C before normalizing the OD_600nm_ of these cultures with carbon-free M17 or MRS, respectively. Cells were then centrifuged (10,000 g / 3 min) and the supernatant was sterile filtered through a 0.22 μm syringe filter. Uninoculated, carbon-free media was also sterile-filtered as a control. Twice-washed *L. plantarum* cells (described above) were resuspended in PBS at an optical density at 600 nm (OD_600nm_) of 1 before inoculation into 96-well culture plates at a final OD_600nm_ of 0.01 in 1:1 CFS (or carbon-free media) to 2x (twice-concentrated) mMRS. The OD_600nm_ was measured over 48 h as described above.

### RNA-seq library construction and transcriptome analysis

The RNA-seq library was constructed and analyzed as previously described (Tejedor-Sanz et al., 2022). In brief, *L. plantarum* NCIMB8826 was grown in mMRS (in triplicate) with or without supplementation of 20 μg/mL DHNA and 1.25 mM ferric ammonium citrate. Cultures were grown at 37 °C to exponential phase (OD_600_ = ∼1.0) before collection via centrifugation (10,000 g / 3 min) at 4 °C. After decanting, cells were flash frozen in liquid N_2_ and stored at -80 °C until RNA extraction with acidic phenol:chloroform:isoamyl alcohol (pH 4.5) as previously described (Golomb et al., 2016). RNA was quantified on a Nanodrop 2000c (ThermoFisher) before two rounds of DNAse digestion using the Turbo DNA-free Kit (Invitrogen, Waltham, MA, USA) following the manufacturer’s protocols. RNA quality was checked using a Bioanalyzer RNA 6000 Nano Kit (Agilent Technologies, Santa Clara, CA, USA) (all RIN values > 9), quantified with the Qubit 2.0 RNA HS Assay (Life Technologies, Carlsbad, CA, USA), and depleted of ribosomal-RNA (rRNA) with the RiboMinus Eukaryote Kit v2 using specific probes for prokaryotic rRNA (ThermoFisher). The remaining RNA was then fragmented to approximately 200 bp, converted to cDNA, and given barcode sequences using the NEBnext Ultra-directional RNA Library Kit for Illumina (New England Biolabs, Ipswitch, MA, USA) with NEBnext Multiplex Oligos for Illumina (Primer Set 1) (New England Biolabs) following the manufacturer’s instructions. The barcoded cDNA libraries were pooled and run across two lanes of a HiSeq400 (Illumina, San Diego, CA, USA) on two separate runs for 150 bp paired-end reads (http://dnatech.genomecenter.ucdavis.edu/).

After sequencing and demultiplexing, DNA sequences for all 12 samples were first visualized in FastQC (ver. 0.11.8) (Andrews, 2010) followed by read trimming with Trimmomatic (ver. 0.39) (Bolger et al., 2014). Remaining reads were aligned to the *L. plantarum* NCIMB8826 chromosome and plasmids using Bowtie2 (ver. 2.3.5) in the [-sensitive] mode (Langmead and Salzberg, 2012) and output “.sam” files were converted to “.bam” files with Samtools (ver 1.9) (Li et al., 2009). Aligned reads which corresponded to NCIMB8826 genes, excluding noncoding sequences (e.g., rRNA, tRNA, trRNA, etc.) were enumerated with FeatureCounts in the [-- stranded=reverse] mode (ver. 1.6.4) (Liao et al., 2014). DESeq2 (Love et al., 2014) using the Wald test in the R-studio shiny app DEBrowser (ver 1.14.2) (Kucukural et al., 2019) was used to quantify differential gene expression based on culture condition. The significance cutoff for differential expression was set to a False-discovery-rate (FDR)-adjusted *p-*value ≤ 0.05 and a Log_2_ (fold-change) ≥ 0.5. Clusters of Orthologous Groups (COGs) were also assigned to genes based the eggNOG (ver. 5.0) database (Huerta-Cepas et al., 2019).

### Hydrogen peroxide production assay

*L. plantarum* NCIMB8826 was grown in mMRS (in triplicate) with or without supplementation of 20 μg/mL DHNA and 1.25 mM ferric ammonium citrate. Cultures were grown at 37 °C to exponential phase (OD_600_ = ∼1.0) before collection via centrifugation (10,000 g / 3 min). Uninoculated cultures (in triplicate) were used for abiotic hydrogen peroxide production and were sampled after 5 h which was when *L. plantarum* cultures reached OD_600_ = 1 in exponential phase. Hydrogen peroxide was measured fluorometrically with the Fluorimetric Hydrogen Peroxide Assay Kit (Sigma-Aldrich) following the manufacturer’s instructions.

### Ferrihydrite reduction assays

*L. plantarum* strains were first incubated in mMRS for 18 h at 37 °C. When indicated, quinones were supplemented at concentrations ranging from 0.01 to 200 μg/mL, and/or ferric ammonium citrate was supplemented at 1.25 mM. Riboflavin was supplemented at 10μM when indicated. Cells were collected via centrifugation (10,000 g, 3 min) and washed twice in PBS. The OD_600nm_ was adjusted to 2 in PBS containing 2.2 mM ferrihydrite (Schwertmann and Fischer, 1973; Stookey, 1970), 2 mM ferrozine (Sigma-Aldrich), and 55 mM mannitol. Quinones and/or riboflavin were supplemented at the above concentrations when indicated, and uninoculated controls were used to subtract background ferrihydrite reduction by quinones. After 3 h incubation at 37 °C, the cells were collected by centrifugation (10,000 g / 5 min) the supernatant was used to determine iron reduction from absorbance measurements at 562 nm with a Synergy 2 microplate reader. Absorbance was converted to the concentration of reduced iron(II) using a standard curve containing a 2-fold range of FeSO_4_ (Sigma-Aldrich) (0.25 mM to 0.016 mM) dissolved in 10 mM cysteine-HCL (RPI, Mount Prospect, IL, USA) and supplemented with 2 mM ferrozine.

In experiments where cell-free supernatant (CFS) was used in place of PBS as the assay medium, overnight cultures of *L. lactis* (GM17) or *Leuconostoc* spp. (gCDM) were grown for 18 h at 30 °C before normalizing the OD_600nm_ of these cultures with carbon-free M17 or CDM, respectively. Cells were then centrifuged (10,000 g / 3 min) and the supernatant was sterile filtered through a 0.22 μm syringe filter. Uninoculated, carbon-free media was also sterile-filtered as a control. The OD_600nm_ of PBS-washed *L. plantarum* cells were then adjusted to 2 in the CFS or uninoculated media which was supplemented with ferrihydrite, ferrozine, and mannitol as described above. Ferrihydrite reduction assays with *L. lactis* or *Leuconostoc* CFS were carried out at 37 °C for 3 h before measuring reduced iron as described above.

### Bioelectrochemical measurements

The bioreactors consisted of double-chamber electrochemical cells (Adams & Chittenden, Berkeley, CA) with a cation exchange membrane (CMI-7000, Membranes International, Ringwood, NJ) that separated them. We used a 3-electrode configuration consisting of an Ag/AgCl (sat. KCl) reference electrode (BASI), a titanium wire counter electrode, and a working electrode of either 6.35-mm-thick graphite felt working electrode of 4x4 cm (Alfa Aesar) with a piece of Ti wire threaded as a current collector and connection to the potentiostat. The bioreactors were sterilized by filling them with ddH2O and autoclaving at 121 °C for 30 min. After this, each chamber media was replaced with 150 mL of filter sterilized CDM media (for the working electrode chamber), and 150 mL of M9 media (BD) (for the counter electrode chamber). The medium (before filter-sterilization) was supplemented with a final concentration of 10 g/L of mannitol and with 20 μg/mL of DHNA diluted 1:1 in DMSO: ddH2O, where appropriate. The media in the working electrode chamber was mixed with a magnetic stir bar for the course of the experiment and N_2_ gas was continuously purged in the working electrode chamber to maintain anaerobic conditions. Four bioreactors were prepared which differed in the CDM used in the working electrode chamber: two bioreactors contained mCDM supplemented with 20 μg/mL of DHNA (diluted 1:1 in DMSO: ddH2O), and other two bioreactors contained mCDM with no DHNA. All the experiments were tested under 30 °C. After approximately 4 h of bubbling the working electrode chamber with N_2_ gas, the working electrode of each bioreactor was polarized. The applied potential to the working electrodes was of 0.2 V versus the Ag/AgCl (sat. KCl) reference electrode. A Bio-Logic Science Instruments potentiostat model VSP-300 was used for performing electrochemical measurements. Once the current density stabilized overnight, the mCDM+DHNA bioreactors were inoculated to a final OD_600nm_ of ∼0.1-0.15 with the cell suspensions of *L. plantarum* prepared in M9 medium. Cell suspensions were prepared from an overnight, statically grown culture (∼16-18 h) of *L. plantarum* cultured in MRS medium at 30 °C. After 45 h of operating the bioreactors and observing current density production, we collected cells from each bioreactor by vigorously shaking the bioreactors to detach cells from the electrode and collecting the medium from the bioreactors (∼150 mL). Cells were collected from each medium by performing 2 cycles of centrifugation (15,228 g, 7 min) and washing with M9 medium. The resulting cell pellets were suspended in oxygen-free M9 medium, and inoculated in the two DHNA-free bioreactors, previously polarized to 0.2 V and left overnight to achieve a stable current density baseline. Cyclic voltammetry analyses were performed at a scan rate of 5 mV/s and in the potential region of -0.7 to 0.4 V vs Ag/AgCl.

### Cellular quinone quantification

To prepare cells for assessments of quinone concentrations, *L. plantarum* was grown in mMRS supplemented with 20 μg/mL DHNA and/or 1.25 mM ferric ammonium citrate (FeAC), *L. lactis* in gM17, and *L. mesenteroides* in gMRS. All strains were incubated for 18 h in their respective culture media at 30 °C prior to collection by centrifugation at 10,000 g for 3 min. Cells were washed twice in PBS before flash freezing with liquid N_2_. The cell pellet was lyophilized for 18 h and then transferred to a 40 mL glass vial, and ground with a spatula and extracted with 3.0 mL of 2:1 dichloromethane (DCM)/MeOH for 2 h while rocking on gently on a shaker at room temperature. The organic solvent was filtered using a glass plug containing celite and dried under vacuum. The crude material was then resuspended in 200 μL of 2:1 isopropanol (IPA)/MeOH. For *L. plantarum* NCIMB8826R incubated in mMRS with DHNA or DHNA and FeAc, data were acquired using an Agilent 6530 LC-q-TOF Mass Spectrometer equipped with an uHPLC system, and data in Supplementary Table S3 were acquired using a Shimadzu 9030 LC-q-ToF Mass Spectrometer equipped with a Nexera LC40 UPLC system. For quinone detection in subsequent experiments (**Table 2**), 5 μL of the crude cell material was analyzed on a LCMS. Menaquinone analogs were quantified using the Phenomenex Luna 5 μm C5 100 Å (50 × 4.6 mm) under the following method: hold 100% solvent A for 5 min then quickly gradient to 80% solvent A/20% solvent B over 0.1 min, then gradient to 100% solvent B over 34.9 min with a flow rate of 0.4 mL/min (solvent A: 95% H2O/5% MeOH +0.1% FA with 5 mM ammonium acetate, solvent B: 60% IPA/35% MeOH/5% H2O + 0.1% FA with 5 mM ammonium acetate). Integrated extracted ion chromatograms for two ion adducts, [M + H]+ and [M+NH4]+, for each menaquinone analog were summed.

### L. plantarum co-culturing experiments

*L. plantarum* NCIMB8826, *L. lactis* TIL46, *L. lactis* TIL999, and *L. mesenteroides* ATCC8293 were grown overnight in gMRS for 18 h at 30 °C. Cells were collected by centrifugation (10,000 g / 3 min) and washed twice in PBS. Approximately 10^7^ CFU/mL of each strain was used to inoculate (in triplicate) 125 mL screw-cap bottles containing gCDM supplemented with 1.25 mM ferric ammonium citrate. Each strain was grown by itself at 30 °C and NCIMB8826R was also grown in co-culture with either TIL46, TIL999, or ATCC8293. At t = 0, 4, 6, 8, and 24 h, cultures were sampled for pH and CFU/mL enumeration on MRS agar (30 °C) for monocultures and mMRS agar (37 °C) for co-cultures to distinguish between *L. plantarum*, *L. lactis*, and *L. mesenteroides* by colony size. At these times, culture aliquots were also centrifuged (10,000 g / 5 min) and the supernatant was collected for Fe^2+^ analysis by immediately adding 2.2 mM ferrozine and using a Fe^2+^ standard curve as described previously.

### Data accession numbers

*L. plantarum* RNA-seq data are available in the NCBI Sequence Read Archive (SRA) under BioProject accession no. PRJNA717240. DEseq2 analysis of RNA-seq data (Supplementary File 1) is available in the Harvard Dataverse: https://doi.org/10.7910/DVN/SAQ5AT

## Acknowledgements

We thank INRAE (National Research Institute for Agriculture, Food – France) and specifically Dr. Véronique Monnet for the provision and use of *Lactococcus lactis* strain TIL46 and TIL999. This work was supported by the National Science Foundation grant #1650042.

## Supplementary Figures

**Supplementary Figure S1.**
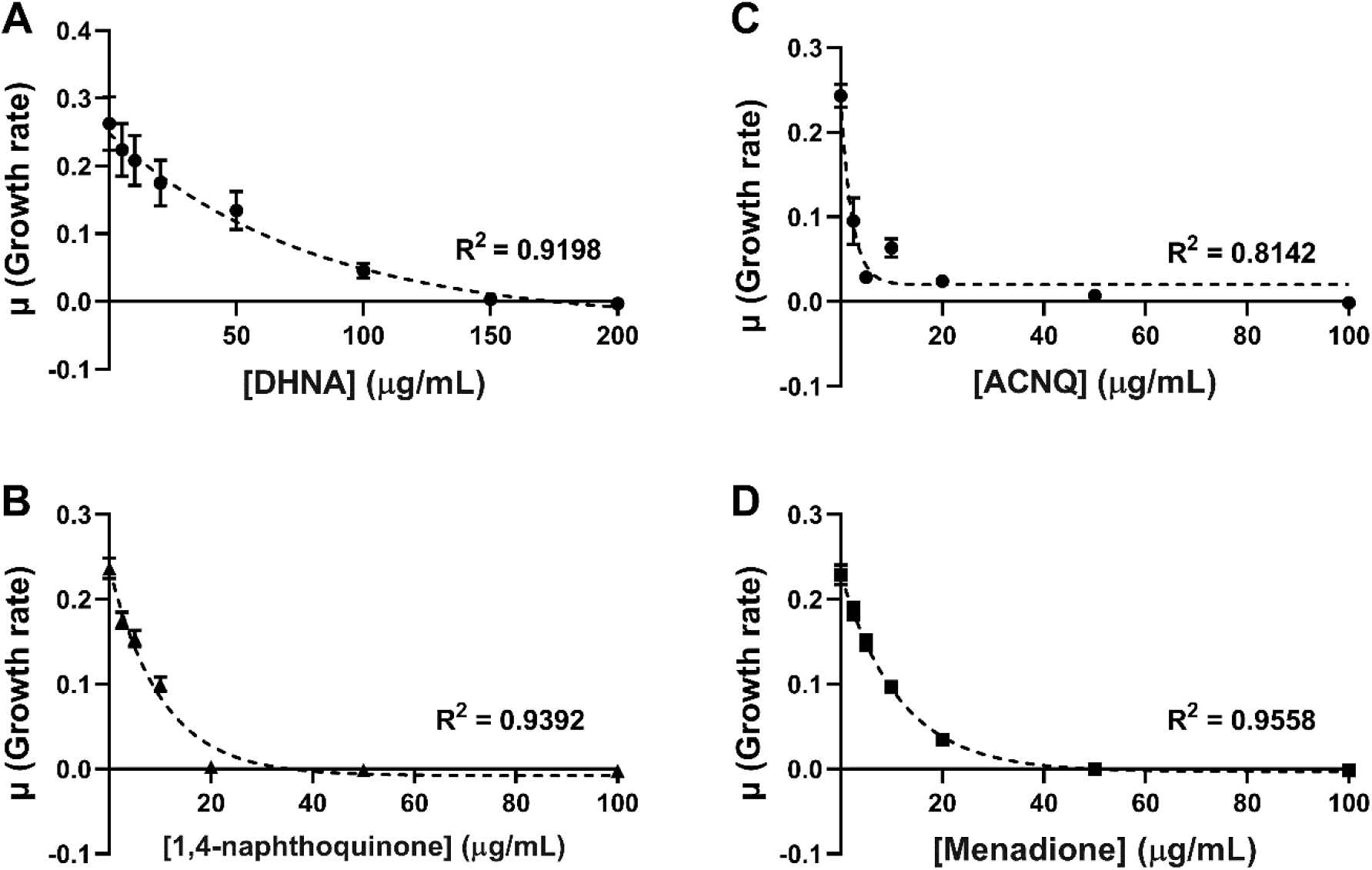
EET-conducive quinones inhibit *L. plantarum* growth. Growth of *L. plantarum* in mMRS supplemented with **(A)** DHNA, **(B)** ACNQ, **(C)** 1,4-naphthoquinone, or **(D)** menadione. The avg ± SEM of three biological replicates is shown.

**Supplementary Figure S2.**
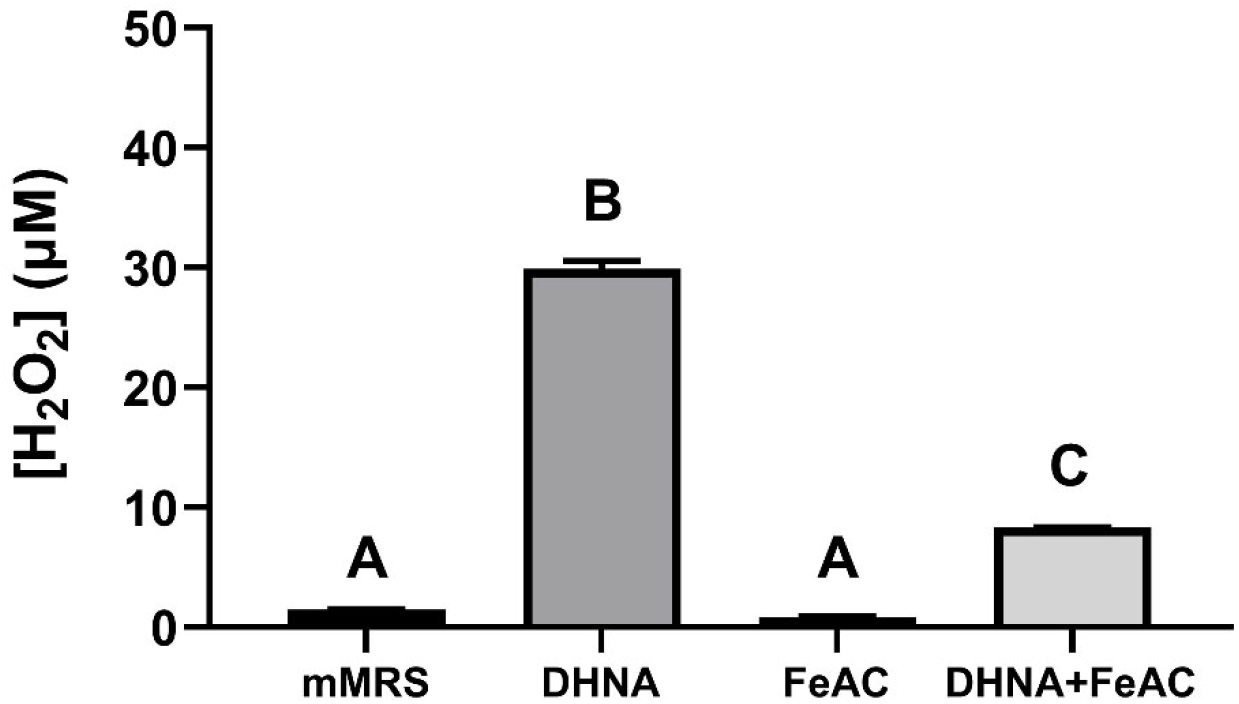
DHNA increases hydrogen peroxide levels in mMRS that are reduced by the presence of FeAC. Hydrogen peroxide produced in mMRS supplemented with 20 μg/mL DHNA and/or 1.25 mM ferric ammonium citrate (Fe<u>A</u>C) after 5 h at 37 °C. *L. plantarum* was inoculated at an OD_600_ = 0.1 and culture supernatant was sampled after 5 h at 37 °C corresponding to when *L. plantarum* reached mid-exponential phase growth. Significant differences in H_2_O_2_ production between culture conditions (represented by letters, p ≤ 0.05) were determined by one-way ANOVA with Tukey’s post-hoc test. The avg ± stdev of three biological replicates is shown.

**Supplementary Figure S3.**
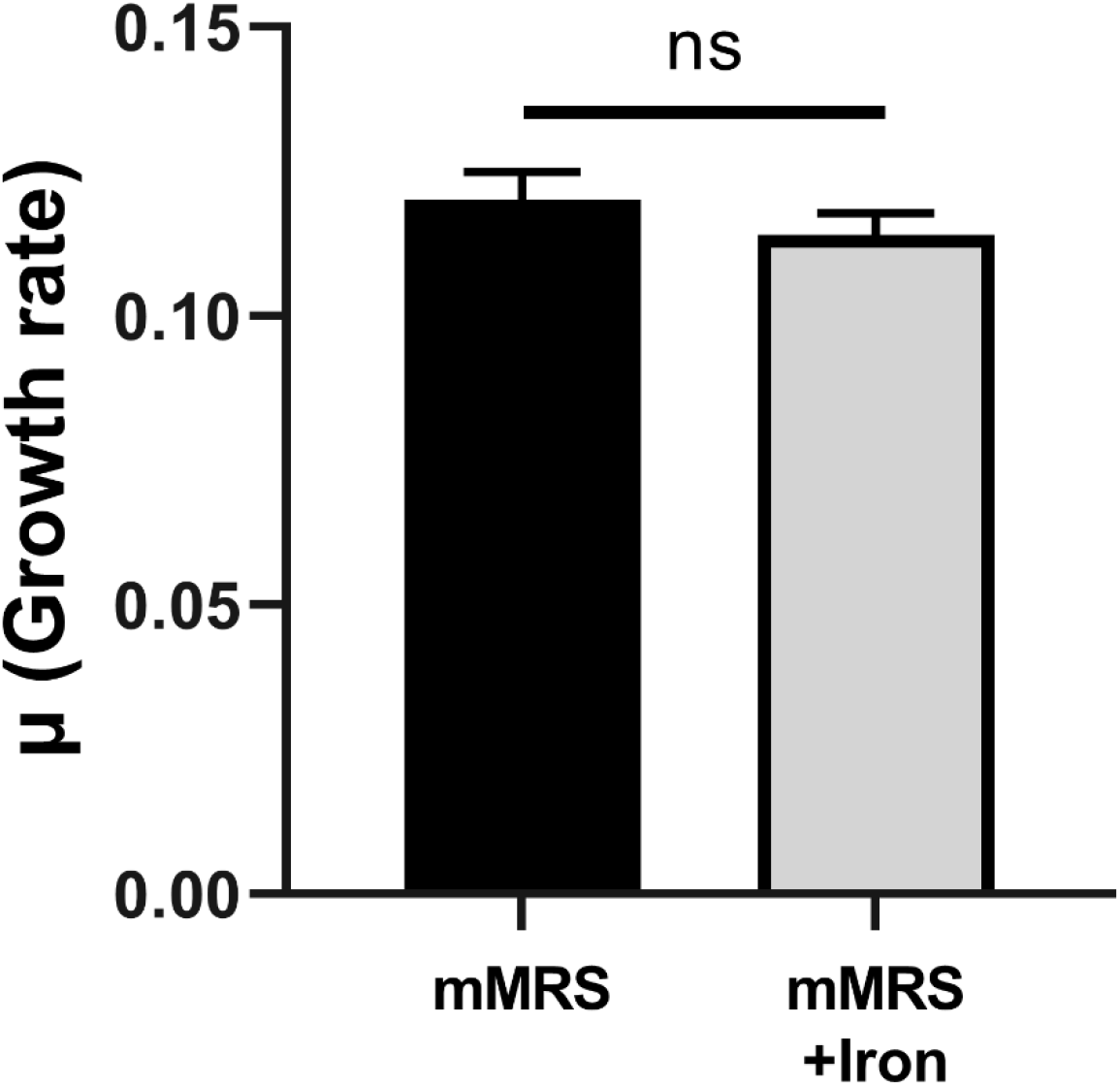
FeAC supplementation in mMRS does not increase *L. plantarum* growth rate. Growth rates were determined by measuring the change in OD_600nm_ per hour during exponential phase. The avg ± SEM of three biological replicates is shown and the absence of statistical difference was determined through a two-tailed t-test.

**Supplementary Figure S4.**
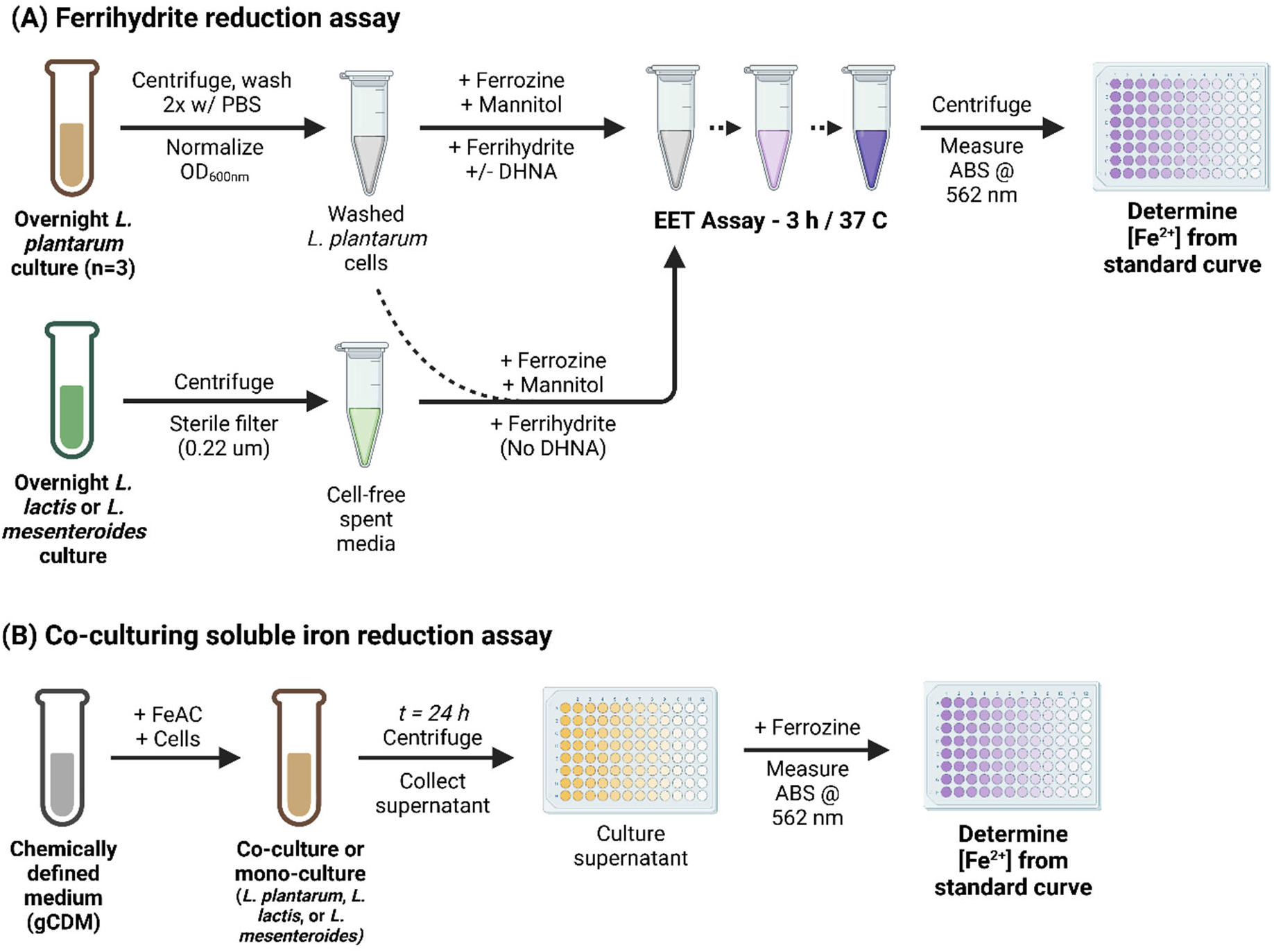
Visualized methods for iron reduction experiments using insoluble or soluble iron. For **(A)** ferrihydrite (insoluble iron), *L. plantarum* cells are collected and washed in 1x PBS (pH 7.2) before normalizing cell numbers via OD_600nm_. EET is performed after ferrozine, mannitol, ferrihydrite, and (where indicated) DHNA or spent growth media from *L. lactis* or *L. mesenteroides* is added. Supernatant is collected after 3 h and used to determine Fe^2+^ from absorbance at 562_nm_. In co-culturing experiments, **(B)** ferric ammonium citrate (soluble iron, FeAC) is added to gCDM along with *L. plantarum*, *L. lactis*, and/or *L. mesenteroides* cells. After 24 h, supernatant is collected and ferrozine is added to instantly determine Fe^2+^ colorimetrically (absorbance at 562_nm_).

**Supplementary Figure S5.**
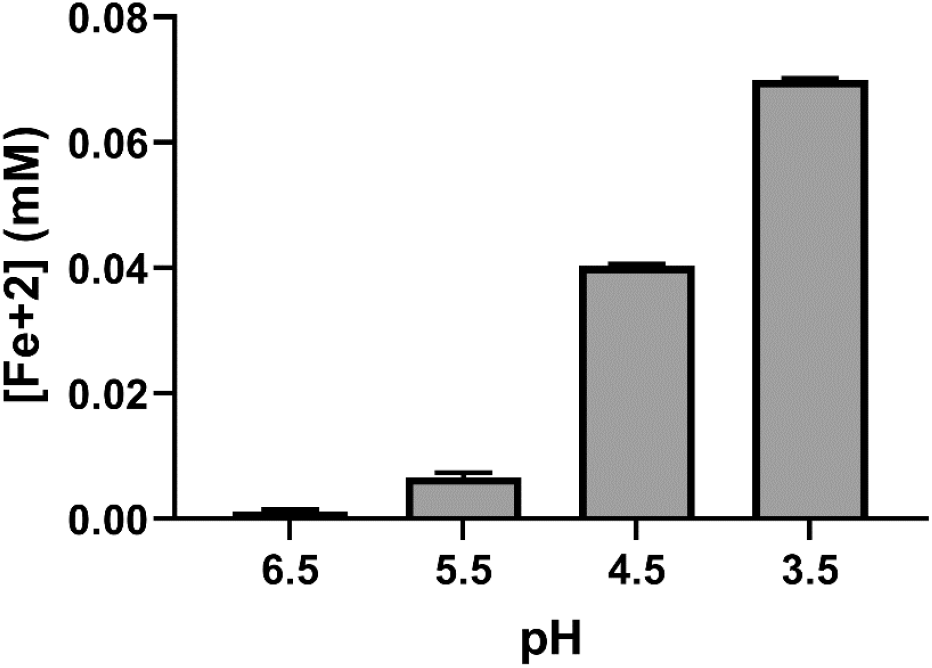
Spontaneous iron reduction in CDM. pH was reduced with ∼4% v/v lactic acid. Iron reduction was detected colorimetrically with 2 mM ferrozine. The avg ± stdev of three replicates is shown.

## Supplementary Tables

**Supplementary Table S1.** Total number of *L. plantarum* genes differentially expressed in DHNA, FeAC, and DHNA+FeAC containing media.

| Growth condition | # Upregulated Genes | # Downregulated Genes | Total # Differentially Expressed Genes |
| --- | --- | --- | --- |
| DHNA | 452 | 473 | 925 |
| FeAC | 35 | 72 | 107 |
| DHNA+FeAC | 123 | 169 | 292 |

Differential expression of all genes in *L. plantarum* during exponential growth in mMRS supplemented with DHNA (20 µg/mL) and/or ferric ammonium citrate (1.25 mM) compared to mMRS. Differences in gene expression levels were considered significant when there was an FDR-adjusted $p$ -value $\leq 0.05$ and $\log_2$ expression fold-change $\geq 0.5$ .

**Supplementary Table S2.**
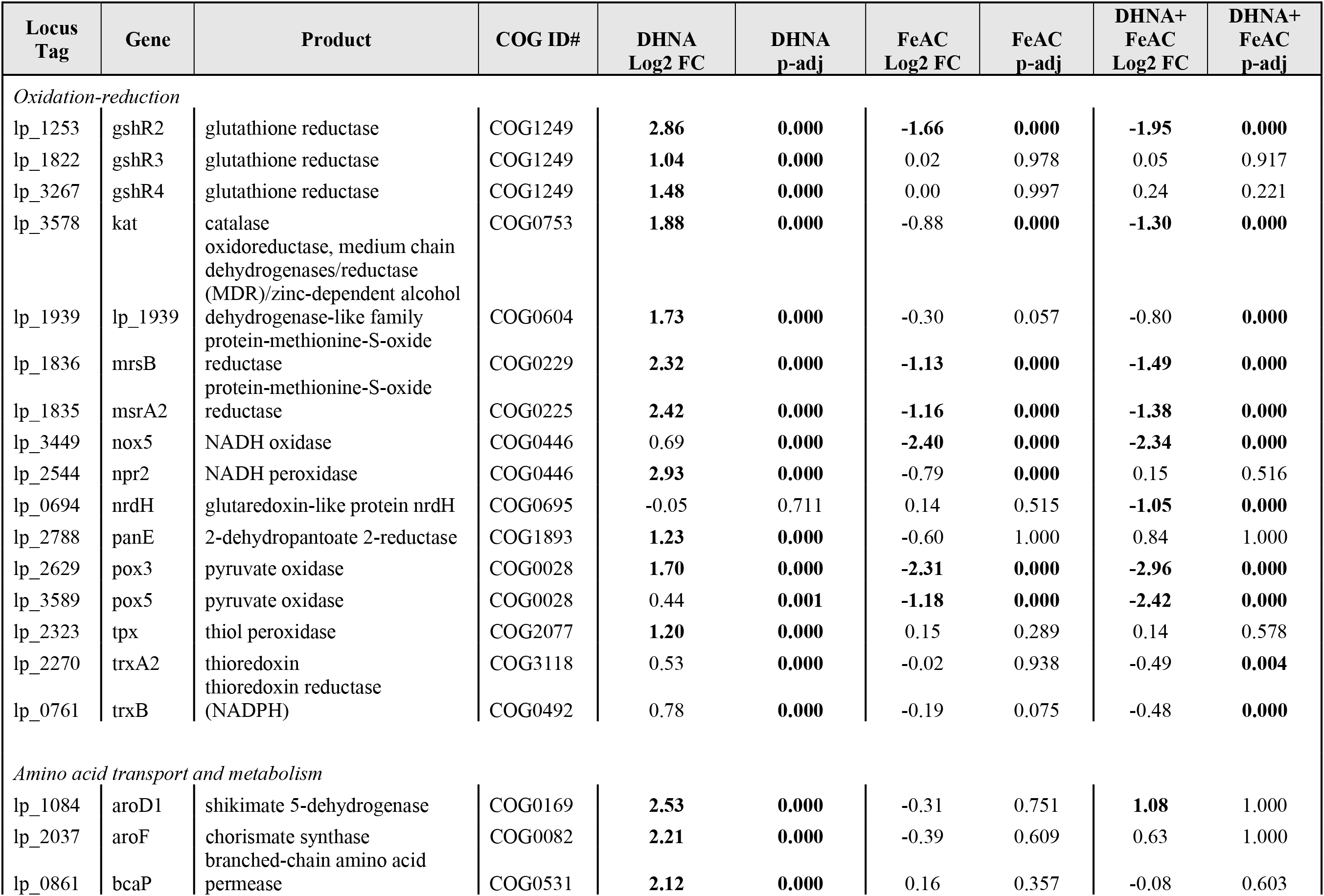
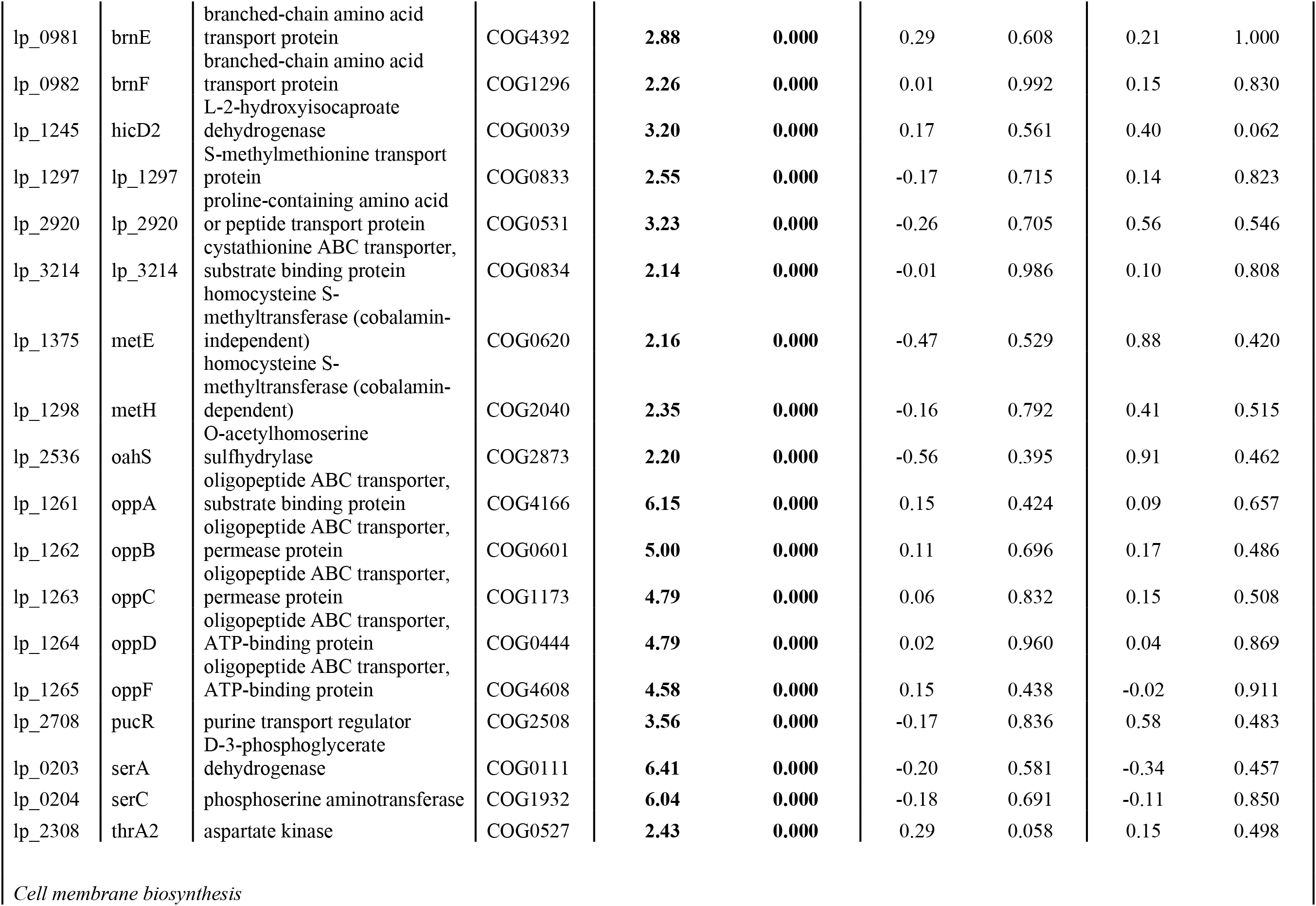
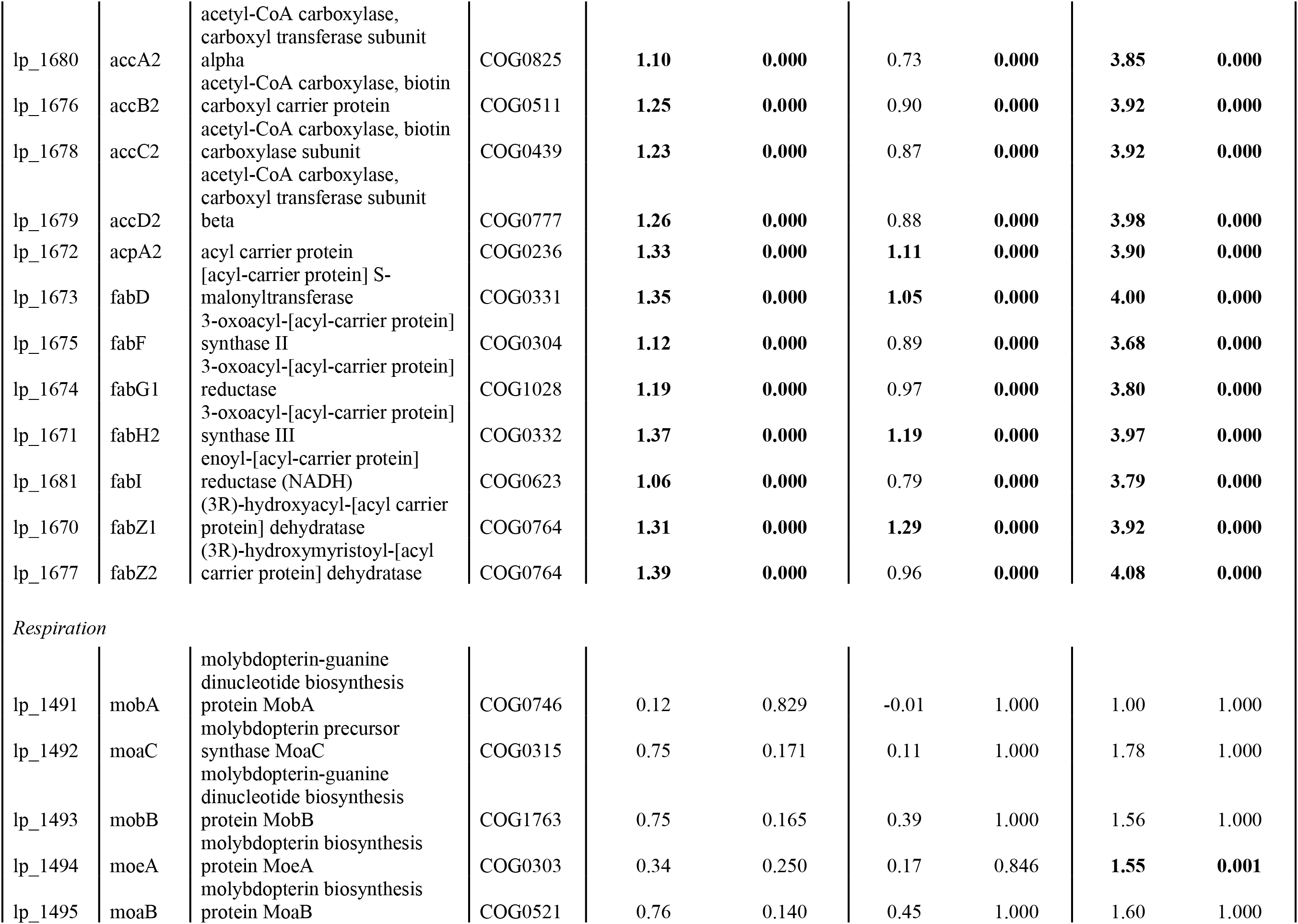
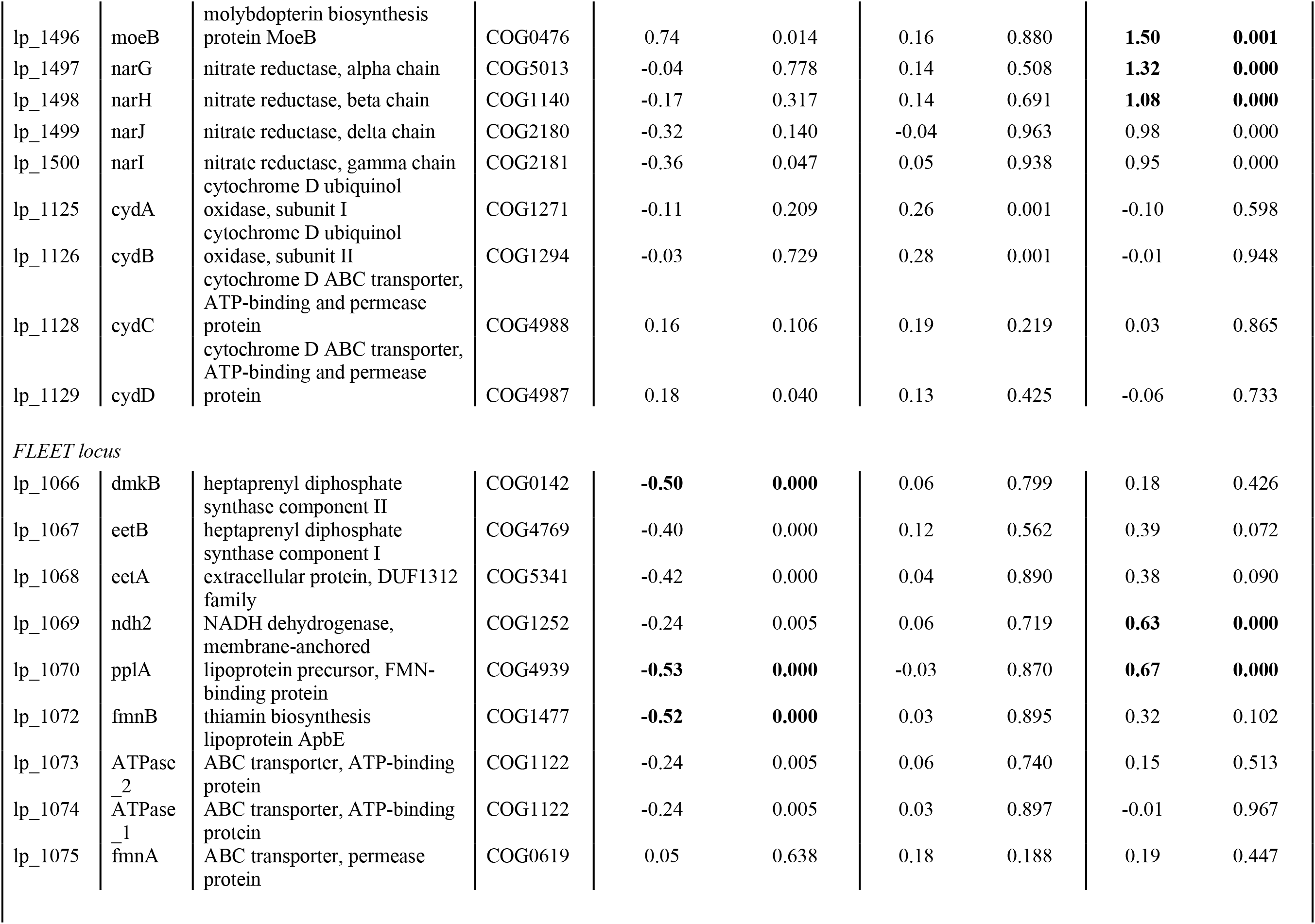
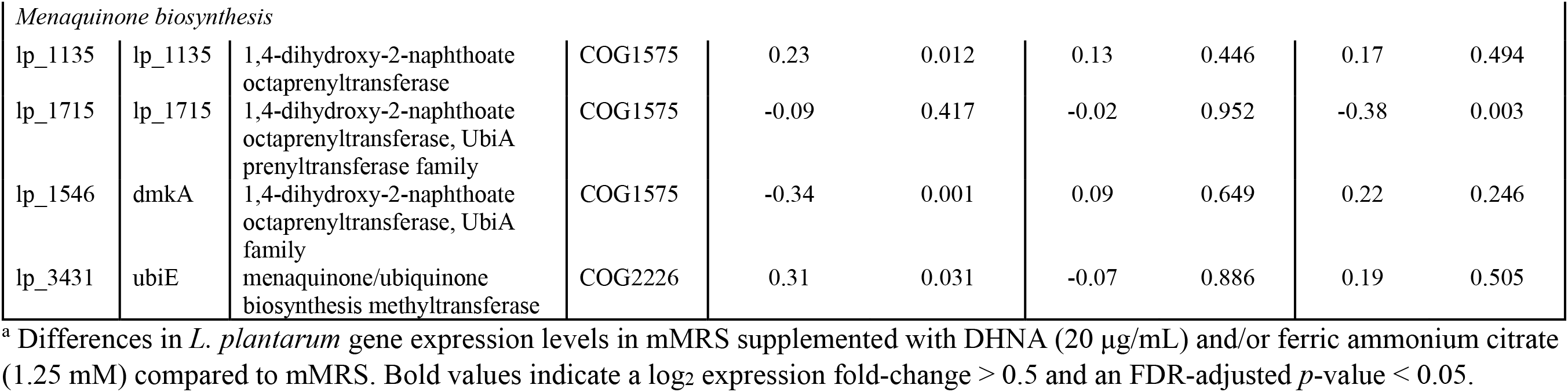
*L. plantarum* NCIMB8826R differentially expressed genes in mMRS with DHNA, FeAC, or DHNA+FeAC.

